# Assessing the conservation and targets of putative sRNAs in *Streptococcus pneumoniae*

**DOI:** 10.1101/2024.11.14.623631

**Authors:** Matthew C. Eichelman, Michelle M. Meyer

**Author notes:** Address correspondence to Michelle M. Meyer,.

## Abstract

RNA regulators are often found in regulatory networks and mediate growth and virulence in bacteria. Small RNAs (sRNAs) are non-coding RNAs that modulate translation initiation and mRNA degradation by base-pairing. To better understand the role of sRNAs in pathogenicity several studies identified sRNAs in *Streptococcus pneumoniae*; however, little functional characterization has followed. This study’s goals are: 1) survey putative sRNAs in *S. pneumoniae*; 2) assess the conservation of these sRNAs; and 3) examine their predicted targets. Three previous studies in *S. pneumoniae* identified 287 putative sRNAs by high-throughput sequencing. This study narrows the candidates to a list of 58 putative sRNAs. BLAST analysis indicates that the 58 sequences are highly conserved across the *S. pneumoniae* pangenome, and 25 of them are identified sporadically in other Streptococcus species. However, only 2 have corresponding sequences identified across several Streptococcus species. We used four RNA-target prediction programs to predict targets for each of the 58 putative sRNAs. Across all probable predictions, six sRNAs have overlapping targets predicted by multiple programs, four of which target numerous transposase encoding transcripts. sRNAs targeting transposase genes display nearly identical and perfect base-pairing. One sRNA, M63 (Spd_sr37), has several probable targets in the CcpA regulon, a network responsible for global catabolite repression, suggesting a possible biological function in carbon metabolism control. Each M63-target interaction exhibits unique base-pairing increasing confidence in the biological relevance of the result. This study produces a list of S*. pneumoniae* putative sRNAs whose predicted targets suggest functional significance in transposon and carbon metabolism regulation.

**Importance:** Previous studies identified many small RNA candidates in *Streptococcus pneumoniae*, several of which were hypothesized to play a role in *S. pneumoniae* virulence. Due to the differing sequencing methods, diverse inclusion criteria, *S. pneumoniae* strain differences, as well as limited follow-up since, it is unclear to what extent candidates identified in different studies have overlapping sequences and functions, and their biological relevance remains ambiguous. This research aims to consolidate the candidate sRNAs across these studies and focuses attention on those that are likely to be regulatory and associated with virulence. This study’s findings enhance our knowledge of the conservation of small regulatory RNAs across the many *Streptococcus pneumoniae* strains and highlights a handful that appear likely to have a role in growth or virulence.

## Introduction

*Streptococcus pneumoniae* is a Gram-positive bacterium that causes various diseases including pneumonia, meningitis, bacteremia, otitis media and sinusitis. Invasive pneumococcal disease is particularly dangerous in children and the elderly (CDC et al., 2013), and in 2004 was responsible for approximately 4 million illness episodes, 445,000 hospitalizations, and 22,000 deaths in the United States (Huang et al., 2011). In 2016, *S. pneumoniae* was the leading cause of lower respiratory infection morbidity and mortality globally causing over a million deaths (GBD 2016 Lower Respiratory Infections Collaborators, 2016). Despite the threat *S. pneumoniae* poses, important components of regulation relating to metabolism and virulence remain less well characterized. Small regulatory RNAs (sRNA) are sequences of 40-500 nucleotides (nt) in length (Li et al., 2012) that can be transcribed by 5’-UTRs, 3’-UTRs, coding, and non-coding sequences (Felden and Augagneur, 2021). However, studies seeking to identify sRNAs tend to focus on intergenic regions because RNAs transcribed in regions lacking an ORF are assumed to be more likely to be functional regulators, whereas those identified within ORFs may be an intermediate RNA decay product from a protein-encoding transcript. Among the different types of sRNAs are *trans*-encoded and *cis*-encoded RNAs. *Trans*-encoded sRNAs regulate genes from distant regions often with imperfect complementarity allowing them to interact with more than one target (Jabbour and Lartigue, 2021). *Cis*-encoded sRNAs act on the mRNA transcript encoded by the opposite DNA strand leading to perfect complementarity (Zorgani et al., 2016) (Fig. 1A). sRNAs modulate the expression of target mRNAs by base pairing to sequester a ribosome binding site or accelerate decay (Papenfort and Vanderpool, 2015). Some sRNAs are dependent on a chaperone like the Hfq or FinO family proteins that have a well-characterized role in aiding the formation of duplexes between sRNAs and their mRNA targets in Enterobacteriaceae. However, the role of RNA chaperones in Gram-positive bacteria is substantially less clear. Such proteins are frequently not present, and when they are present their functionality is often substantially different from that observed in gram-negative organisms (Christopoulou and Granneman, 2022). In *S. pneumoniae*, recent work shows KH domain proteins, like KhpA and KhpB, are associated with sRNAs (Olejiiczak et al., 2022), and sRNAs have been associated with the exonuclease Cbf1 (Hor et al. 2020). However, there are no confirmed sRNA chaperone homologs in the *S. pneumoniae* genome (Zhang et al., 2003).

**Figure 1:**
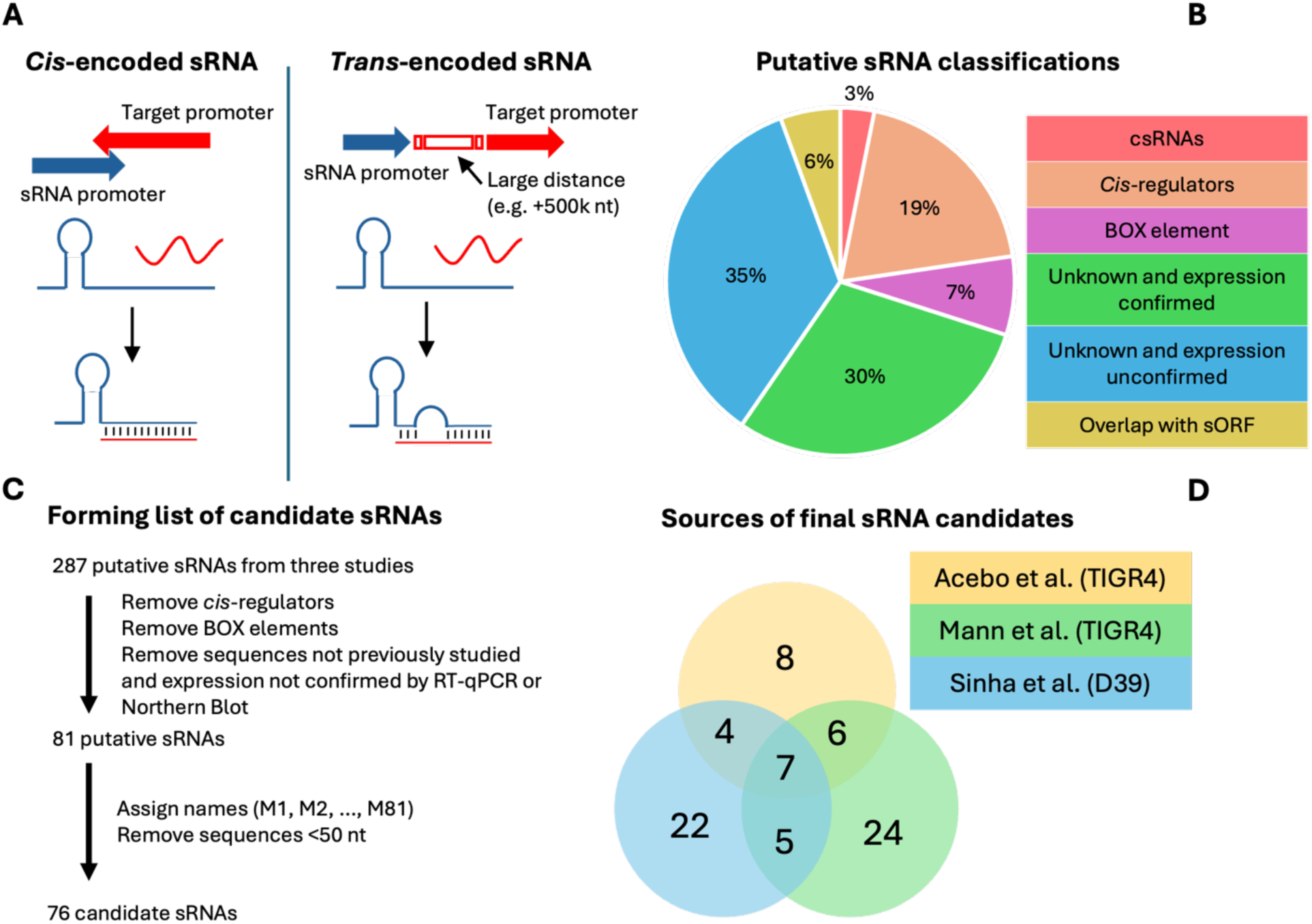
**A**) The *cis*-encoded sRNA, found on the strand opposite to the mRNA coding strand, binds the target mRNA with perfect complementarity. The *trans*-encoded sRNA, expressed from a region distant to the target, binds with imperfect complementarity. **B)** The classifications of the 287 putative sRNAs. **C)** Process of narrowing the 287 putative sRNAs to a list of 76 candidates for further analysis. **D)** Among the candidates, 7 were identified by all three studies and 15 by two of the three studies.

Identification of putative sRNAs in *S. pneumoniae* has been performed several times, but follow-up characterization has been limited. The exception is a group of sRNAs called cia-dependent sRNAs (csRNAs) that have been studied across Streptococcus with considerable work done in *S. pneumoniae*. The csRNAs are controlled by the CiaRH two-component system (TCS) that is involved in natural competence and general virulence (Halfmann et al., 2007, Marx et al., 2010, Tsui et al., 2010). In *S. pneumoniae* the CiaRH TCS expresses five sRNAs, with experimentally verified targets, that prevent autolysis triggered by various conditions, like the presence of deoxycholate, to allow the maintenance of stationary growth phase (Mascher et al., 2006). The csRNAs have also recently been implicated in promoting Zn homeostasis (De Lay et al., 2024). Three previous studies identified hundreds of additional putative sRNAs using diverse inclusion criteria. Some of these sequences may be annotated as homologs of RNA families such as Pyr elements (RF00515) or TPP riboswitches (RF00059), BOX elements (AT-rich repeats that are highly transcribed), or ribosomal protein leaders (sequences in the 5’-UTR of ribosomal protein transcripts that control the concentration of the ribosomal protein) (Mann et al., 2012, Fu et al., 2013, Babina et al., 2015). However, it remains unclear which of the remaining sequences are regulatory. Here we find that 58 of the putative sRNAs are highly conserved in the *S. pneumoniae* pangenome and we predict the mRNA targets of these sRNAs. The predictions suggest 7 putative sRNAs, which have highly probable targets, interact with mRNAs coding for transposases and genes involved in carbon metabolism regulation.

## Results and Discussion

### *S. pneumoniae* genome contains +70 putative sRNAs

In assessing which previously identified sRNA candidates are likely to have a biological function and prioritize candidates for further investigation we examined a pool of 287 putative sRNAs originating from three studies (Acebo et al., 2012, Mann et al., 2012, and Sinha et al., 2019, Additional Datafile 1). We note 65% are functionally uncharacterized whereas the other 35% may be annotated as homologs of *cis*-regulators, BOX elements, csRNAs, or overlap with small open reading frames (sORF) (Fig. 1B). We further narrowed this pool to a list where each sRNA has at least 1 of 3 attributes: 1) sequence identified in multiple studies, 2) expression confirmed by Northern blot or RT-qPCR, 3) sequence characterized as a csRNA or overlapping with a sORF (Laczkovich et al., 2022). *Cis*-regulators and BOX elements were also excluded from our list. Thus, the new list contains 81 putative sRNAs that were assigned names “M1” through “M81”. However, 5 sequences are <50 nt and were subsequently removed from the list leaving a final total of 76 candidate sRNAs (Fig. 1C). We observe that many of these sRNAs were found in a single study emphasizing the different sequencing strategies and inclusion criteria of the previous studies (Fig. 1D). After compiling the final list for further analysis, we conclude there are over 70 putative sRNAs in the *S. pneumoniae* genome (Additional Datafile 2).

### Majority of sRNA candidates are conserved across the *S. pneumoniae* pangenome

To increase our confidence in the biological relevance of the putative sRNAs and prioritize them for further investigation, we assessed the conservation of the candidate sRNAs across the genomes of 385 *S. pneumoniae* strains. BLAST (Altschul et al., 1990) analysis indicated 70/76 candidates are present in the genomes for a majority of 385 *S. pneumoniae* strains (Fig. 2A) (Cremers et al. 2015, Rosconi et al., 2022). Among these 70, only 60 candidates appear to be non-repetitive sequences, all of which display average sequence identity >97% to the best hit in each genome, indicating the sequences are highly conserved across the *S. pneumoniae* pangenome (Table 1). Interestingly, all of the 6 sRNAs found in <12 strains were identified in the *S. pneumoniae* strain TIGR4 (Fig. 2), highlighting the possibility for strain specific sRNAs in *S. pneumoniae*. Two of the final candidates, M8 and M12, were identified as *cis*-regulatory elements later during our analysis and removed, leaving a total of 58 sRNAs for further analysis. From this final group of 58 candidates, we observed that synteny is preserved (see Methods) across the *S. pneumoniae* strains in 43 of the 58 final candidates. To determine if any of the sRNAs are conserved in species other than *S. pneumoniae* we also analyzed a range of other species in the Streptococcus genus. Only 2 of the candidate sRNAs (M77 and M81) align with sequence identities >65% to each of *S. pyogenes*, *S. mutans*, *S. suis*, *S. mitis*, *S. oralis*, and *S. gordonii,* and 25 additional sRNA candidates can be identified in a subset of these organisms (Supplemental Table S1). Thus, it appears over half of the sRNAs are unique to *S. pneumoniae*, consistent with the narrow distribution of many sRNAs across other bacterial species (Peer and Margalit, 2014).

**Figure 2:**
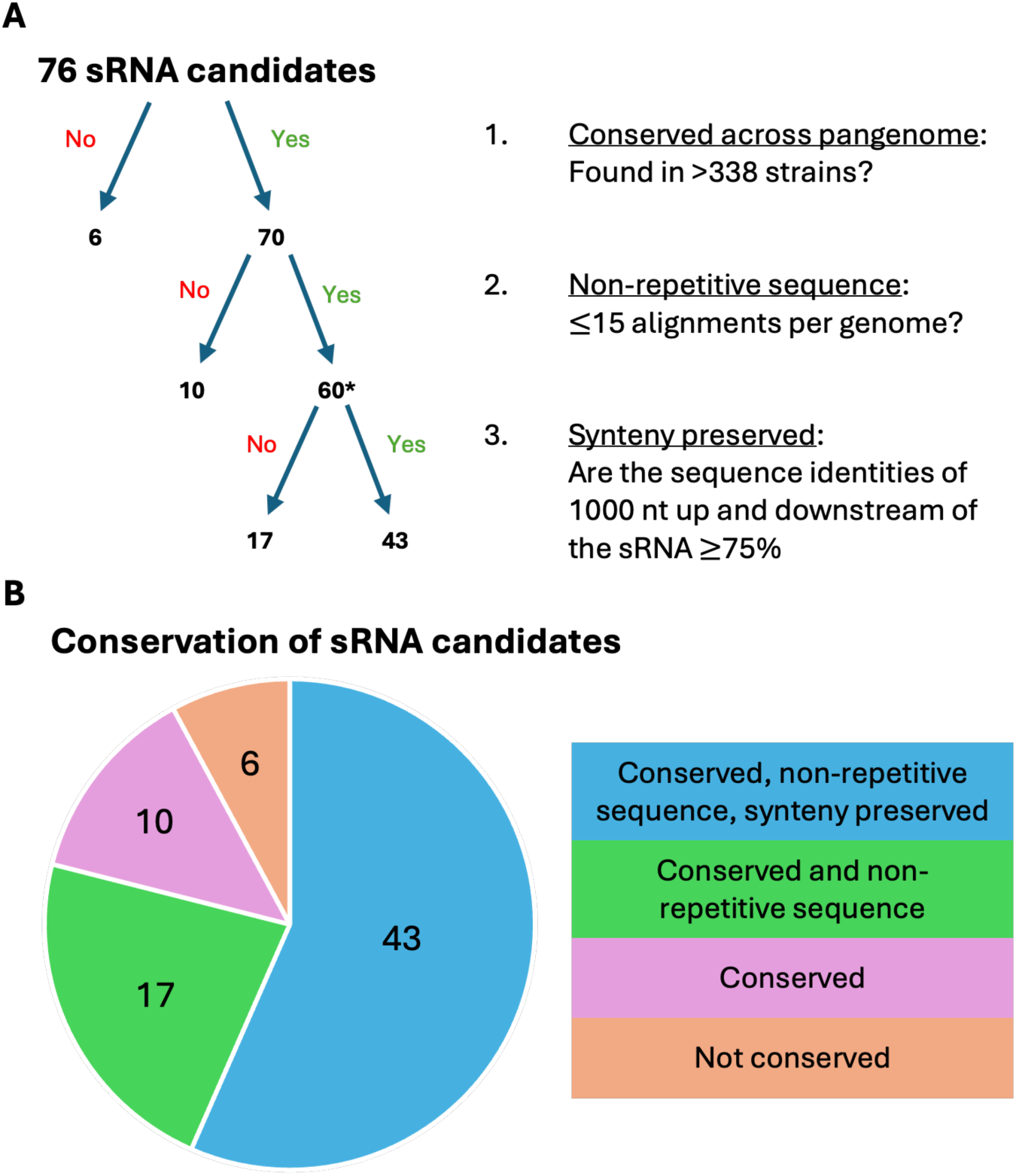
**A)** The conservation of sRNA candidates across the *S. pneumoniae* pangenome. The * next to “60” highlights that 2 sRNAs were later removed to form the final list of 58 sRNAs (see Table 1 legend). **B)** The four degrees of conservation of the sRNA candidates. “Conserved”, “non-repetitive sequence”, and “synteny preserved” refer to the criteria for each level of the tree diagram in part A.

**Table 1:**
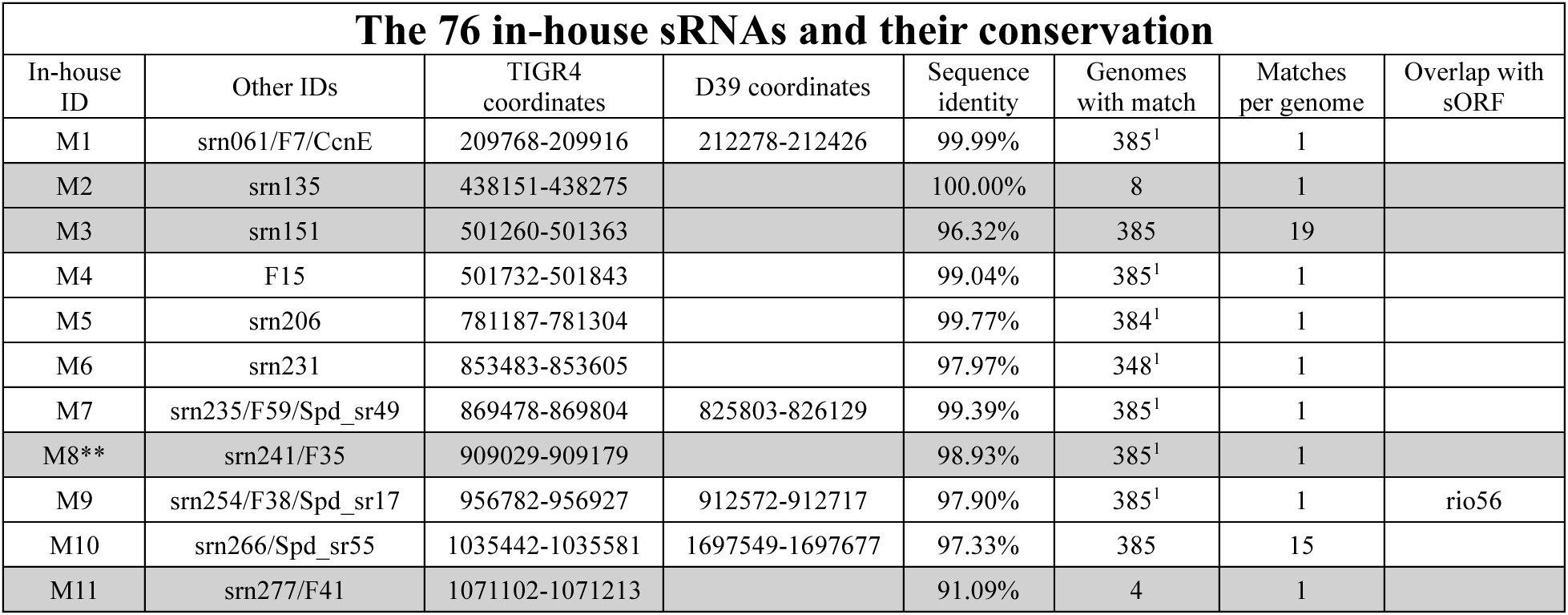
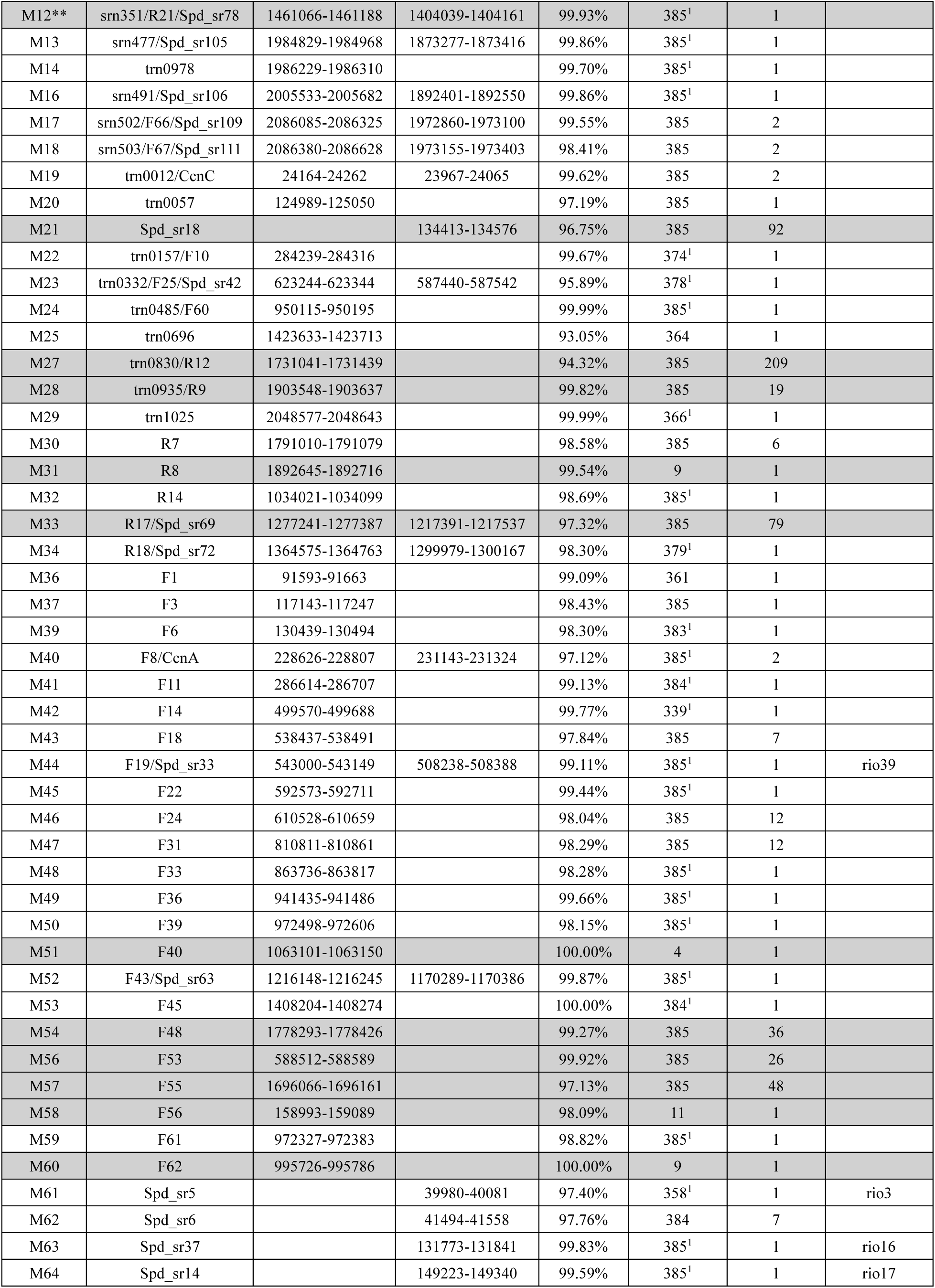
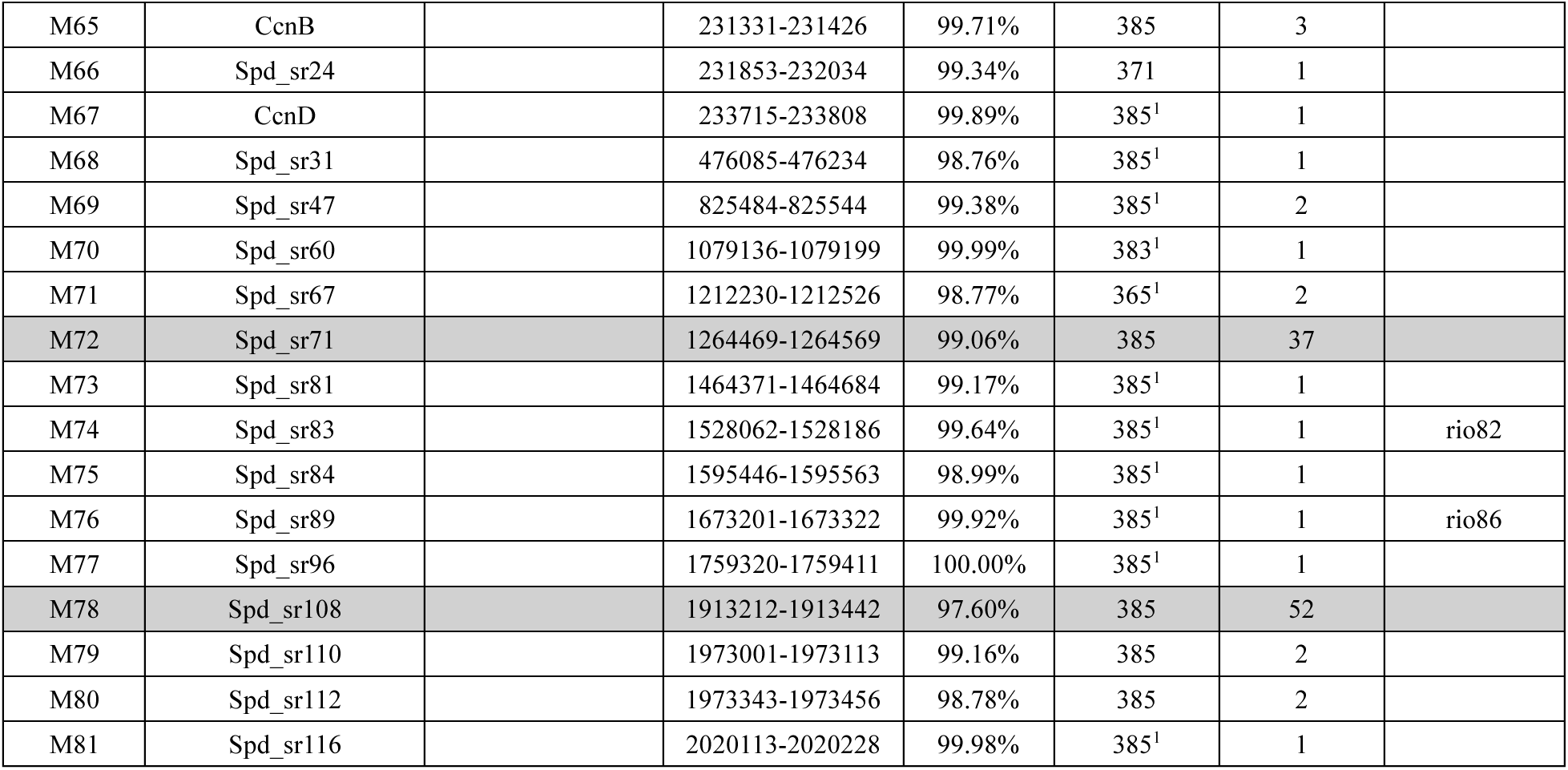
The 76 candidate sRNAs and their conservation. In the “Other IDs” column, sRNAs with the prefix “srn”, “trn”, “R”, and “F” were identified in TIGR4 (Acebo et al., 2012, Mann et al., 2012) and sRNAs with the prefix “Spd_sr” and “Ccn” were identified in D39W (Sinha et al., 2019). The TIGR4 coordinates indicate the sequence location in the NC_003028.3 genome. Likewise, D39 coordinates corresponds to the NC_008533.2 genome. The “Sequence identity” and “Matches per genome” are averages across the 385 strains. Preservation of synteny indicated by ^1^ in the “Genomes with match” column. **M8 later identified as a *cis*-regulator (pyrR element: RFAM:RF00515), and M12 identified as a Mn^2+^ responsive riboswitch (Martin et al. 2019, RFAM:RF000080). The “Overlap with sORF” column indicates the sRNA overlaps with a previously identified sORF (Laczkovich et al., 2022). White rows correspond to the 58 final sRNAs that are conserved in many *S. pneumoniae strains*, but have fewer than 15 instances per genome.

| The 76 in-house sRNAs and their conservation |  |  |  |  |  |  |  |
| --- | --- | --- | --- | --- | --- | --- | --- |
| In-house ID | Other IDs | TIGR4 coordinates | D39 coordinates | Sequence identity | Genomes with match | Matches per genome | Overlap with sORF |
| M1 | srn061/F7/CcnE | 209768-209916 | 212278-212426 | 99.99% | 385<sup>1</sup> | 1 |  |
| M2 | srn135 | 438151-438275 |  | 100.00% | 8 | 1 |  |
| M3 | srn151 | 501260-501363 |  | 96.32% | 385 | 19 |  |
| M4 | F15 | 501732-501843 |  | 99.04% | 385<sup>1</sup> | 1 |  |
| M5 | srn206 | 781187-781304 |  | 99.77% | 384<sup>1</sup> | 1 |  |
| M6 | srn231 | 853483-853605 |  | 97.97% | 348<sup>1</sup> | 1 |  |
| M7 | srn235/F59/Spd_sr49 | 869478-869804 | 825803-826129 | 99.39% | 385<sup>1</sup> | 1 |  |
| M8** | srn241/F35 | 909029-909179 |  | 98.93% | 385<sup>1</sup> | 1 |  |
| M9 | srn254/F38/Spd_sr17 | 956782-956927 | 912572-912717 | 97.90% | 385<sup>1</sup> | 1 | rio56 |
| M10 | srn266/Spd_sr55 | 1035442-1035581 | 1697549-1697677 | 97.33% | 385 | 15 |  |
| M11 | srn277/F41 | 1071102-1071213 |  | 91.09% | 4 | 1 |  |
| M12** | srn351/R21/Spd_sr78 | 1461066-1461188 | 1404039-1404161 | 99.93% | 385<sup>1</sup> | 1 |  |
| M13 | srn477/Spd_sr105 | 1984829-1984968 | 1873277-1873416 | 99.86% | 385<sup>1</sup> | 1 |  |
| M14 | trn0978 | 1986229-1986310 |  | 99.70% | 385<sup>1</sup> | 1 |  |
| M16 | srn491/Spd_sr106 | 2005533-2005682 | 1892401-1892550 | 99.86% | 385<sup>1</sup> | 1 |  |
| M17 | srn502/F66/Spd_sr109 | 2086085-2086325 | 1972860-1973100 | 99.55% | 385 | 2 |  |
| M18 | srn503/F67/Spd_sr111 | 2086380-2086628 | 1973155-1973403 | 98.41% | 385 | 2 |  |
| M19 | trn0012/CcnC | 24164-24262 | 23967-24065 | 99.62% | 385 | 2 |  |
| M20 | trn0057 | 124989-125050 |  | 97.19% | 385 | 1 |  |
| M21 | Spd_sr18 |  | 134413-134576 | 96.75% | 385 | 92 |  |
| M22 | trn0157/F10 | 284239-284316 |  | 99.67% | 374<sup>1</sup> | 1 |  |
| M23 | trn0332/F25/Spd_sr42 | 623244-623344 | 587440-587542 | 95.89% | 378<sup>1</sup> | 1 |  |
| M24 | trn0485/F60 | 950115-950195 |  | 99.99% | 385<sup>1</sup> | 1 |  |
| M25 | trn0696 | 1423633-1423713 |  | 93.05% | 364 | 1 |  |
| M27 | trn0830/R12 | 1731041-1731439 |  | 94.32% | 385 | 209 |  |
| M28 | trn0935/R9 | 1903548-1903637 |  | 99.82% | 385 | 19 |  |
| M29 | trn1025 | 2048577-2048643 |  | 99.99% | 366<sup>1</sup> | 1 |  |
| M30 | R7 | 1791010-1791079 |  | 98.58% | 385 | 6 |  |
| M31 | R8 | 1892645-1892716 |  | 99.54% | 9 | 1 |  |
| M32 | R14 | 1034021-1034099 |  | 98.69% | 385<sup>1</sup> | 1 |  |
| M33 | R17/Spd_sr69 | 1277241-1277387 | 1217391-1217537 | 97.32% | 385 | 79 |  |
| M34 | R18/Spd_sr72 | 1364575-1364763 | 1299979-1300167 | 98.30% | 379<sup>1</sup> | 1 |  |
| M36 | F1 | 91593-91663 |  | 99.09% | 361 | 1 |  |
| M37 | F3 | 117143-117247 |  | 98.43% | 385 | 1 |  |
| M39 | F6 | 130439-130494 |  | 98.30% | 383<sup>1</sup> | 1 |  |
| M40 | F8/CcnA | 228626-228807 | 231143-231324 | 97.12% | 385<sup>1</sup> | 2 |  |
| M41 | F11 | 286614-286707 |  | 99.13% | 384<sup>1</sup> | 1 |  |
| M42 | F14 | 499570-499688 |  | 99.77% | 339<sup>1</sup> | 1 |  |
| M43 | F18 | 538437-538491 |  | 97.84% | 385 | 7 |  |
| M44 | F19/Spd_sr33 | 543000-543149 | 508238-508388 | 99.11% | 385<sup>1</sup> | 1 | rio39 |
| M45 | F22 | 592573-592711 |  | 99.44% | 385<sup>1</sup> | 1 |  |
| M46 | F24 | 610528-610659 |  | 98.04% | 385 | 12 |  |
| M47 | F31 | 810811-810861 |  | 98.29% | 385 | 12 |  |
| M48 | F33 | 863736-863817 |  | 98.28% | 385<sup>1</sup> | 1 |  |
| M49 | F36 | 941435-941486 |  | 99.66% | 385<sup>1</sup> | 1 |  |
| M50 | F39 | 972498-972606 |  | 98.15% | 385<sup>1</sup> | 1 |  |
| M51 | F40 | 1063101-1063150 |  | 100.00% | 4 | 1 |  |
| M52 | F43/Spd_sr63 | 1216148-1216245 | 1170289-1170386 | 99.87% | 385<sup>1</sup> | 1 |  |
| M53 | F45 | 1408204-1408274 |  | 100.00% | 384<sup>1</sup> | 1 |  |
| M54 | F48 | 1778293-1778426 |  | 99.27% | 385 | 36 |  |
| M56 | F53 | 588512-588589 |  | 99.92% | 385 | 26 |  |
| M57 | F55 | 1696066-1696161 |  | 97.13% | 385 | 48 |  |
| M58 | F56 | 158993-159089 |  | 98.09% | 11 | 1 |  |
| M59 | F61 | 972327-972383 |  | 98.82% | 385<sup>1</sup> | 1 |  |
| M60 | F62 | 995726-995786 |  | 100.00% | 9 | 1 |  |
| M61 | Spd_sr5 |  | 39980-40081 | 97.40% | 358<sup>1</sup> | 1 | rio3 |
| M62 | Spd_sr6 |  | 41494-41558 | 97.76% | 384 | 7 |  |
| M63 | Spd_sr37 |  | 131773-131841 | 99.83% | 385<sup>1</sup> | 1 | rio16 |
| M64 | Spd_sr14 |  | 149223-149340 | 99.59% | 385<sup>1</sup> | 1 | rio17 |
| M65 | CcnB |  | 231331-231426 | 99.71% | 385 | 3 |  |
| M66 | Spd_sr24 |  | 231853-232034 | 99.34% | 371 | 1 |  |
| M67 | CcnD |  | 233715-233808 | 99.89% | 385<sup>1</sup> | 1 |  |
| M68 | Spd_sr31 |  | 476085-476234 | 98.76% | 385<sup>1</sup> | 1 |  |
| M69 | Spd_sr47 |  | 825484-825544 | 99.38% | 385<sup>1</sup> | 2 |  |
| M70 | Spd_sr60 |  | 1079136-1079199 | 99.99% | 383<sup>1</sup> | 1 |  |
| M71 | Spd_sr67 |  | 1212230-1212526 | 98.77% | 365<sup>1</sup> | 2 |  |
| M72 | Spd_sr71 |  | 1264469-1264569 | 99.06% | 385 | 37 |  |
| M73 | Spd_sr81 |  | 1464371-1464684 | 99.17% | 385<sup>1</sup> | 1 |  |
| M74 | Spd_sr83 |  | 1528062-1528186 | 99.64% | 385<sup>1</sup> | 1 | rio82 |
| M75 | Spd_sr84 |  | 1595446-1595563 | 98.99% | 385<sup>1</sup> | 1 |  |
| M76 | Spd_sr89 |  | 1673201-1673322 | 99.92% | 385<sup>1</sup> | 1 | rio86 |
| M77 | Spd_sr96 |  | 1759320-1759411 | 100.00% | 385<sup>1</sup> | 1 |  |
| M78 | Spd_sr108 |  | 1913212-1913442 | 97.60% | 385 | 52 |  |
| M79 | Spd_sr110 |  | 1973001-1973113 | 99.16% | 385 | 2 |  |
| M80 | Spd_sr112 |  | 1973343-1973456 | 98.78% | 385 | 2 |  |
| M81 | Spd_sr116 |  | 2020113-2020228 | 99.98% | 385<sup>1</sup> | 1 |  |

### Several sRNA candidates overlap with reported sORFs

Of the 58 final sRNAs candidates, 6 of them overlap with previously identified sORFs (Table 1) (Laczkovich et al., 2022), and one (M64) is complementary to an sORF. Moreover, we see that these 5 of the 7 sORFs are found in nearly all 385 strains with 1 alignment per genome and a high degree of conservation with nucleotide and peptide sequence identities >98%. The first exception, rio56, is a 6 amino acid peptide sequence and is too short for BLAST to produce alignments with an e-value < 20, and the second, rio86, has an average peptide sequence identity of 93.6%. Of particular interest is M61 (Spd_sr5/srf-02), which overlaps with rio3, whose expression was shown to promote *in vivo* fitness (Laczkovich et al., 2022). However, dual-function RNAs, sequences with coding and non-coding functions, are known to exist in other bacteria like SgrT/SgrS and AzuC/AzuR in *Escherichia coli* (Aoyama and Storz, 2023) and Pel RNA in *Streptococcus pyogenes* (Raina et al., 2018). Thus, we decided to retain sORF overlapping sRNAs for target analysis.

### RNA target prediction programs struggle to correctly predict validated targets

Several RNA-RNA interaction prediction (RIP) programs have been developed to predict mRNA:sRNA interactions with newer models displaying the highest accuracies. These include IntaRNA (Busch et al., 2008, Wright et al., 2014, Mann et al., 2017, Raden et al., 2018), CopraRNA (Wright et al., 2013, Wright et al., 2014, Raden et al., 2018), sRNARFTarget (Naskulwar and Peña-Castillo, 2022), and TargetRNA3 (Tjaden, 2023). The programs take various approaches with newer RIP programs implementing machine learning algorithms.

Despite the improvement over time, all the tools have a high false positive rate (Tjaden, 2023). Moreover, most of the data on which the models are validated and trained is from Hfq-dependent sRNA networks in *E. coli* that may not be reflective of sRNA-target interactions in organisms without Hfq like *S. pneumoniae*. This poses a challenge to determining the validity of a predicted target through computational methods alone. By examining the targets of multiple programs with different approaches we hope to increase the confidence in the validity of predicted sRNA-target pairs.

As a baseline evaluation of the RIP programs, we compared the predicted and known targets of the csRNAs using IntaRNA, sRNARFTarget, and TargetRNA3 (Schnorpfeil et al., 2013). None of the programs correctly predicted any of the known csRNA targets (SP_2237/SP_RS11435, SP_0090/ SP_RS00460, SP_0161/ SP_RS00830, SP_0626/SP_RS03070, and SP_1215/SP_RS05965) as the most likely target. If we include the top five most likely targets, then IntaRNA correctly predicts that csRNAs 2 and 3 target SP_RS00460 and both IntaRNA and TargetRNA3 correctly predicted that csRNA4 targets SP_RS00460. We also confirmed the sequences that we examined are consistent with reported 5’ and 3’ RNA-seq data in TIGR4 (Warrier et al., 2018, Furumo and Meyer, 2024) to ensure our inputs were not causing the low accuracy. These results support the existing notion that even the best RIP programs suffer from a high false positive rate, but do provide informative results.

### RIP programs predict thousands of sRNA-target pairs

We used multiple programs to make target predictions for the candidates. For all 58 sRNAs we used IntaRNA, sRNARFTarget, and TargetRNA3. Targets were predicted in six different *S. pneumoniae* strains: TIGR4, D39, and four arbitrarily selected strains from PRJNA514780 (Rosconi et al., 2022) (see Methods). We also used CopraRNA, but only for the 3 sRNAs with sequence identities >65% in at least 4 related Streptococcus species due to the algorithm’s comparative approach to target identification. CopraRNA was used to make predictions for *S. pneumoniae*, *S. pyogenes*, *S. mutans*, *S. suis*, *S. mitis*, *S. oralis*, and *S. gordonii*. Each program produces a variable number of outputs per sRNA per strain/species. IntaRNA and CopraRNA made five predictions (a customizable parameter), sRNARFTarget predicted a probability for every gene in the *S. pneumoniae* transcriptome (>2000 genes), and TargetRNA3 reported a variable number of targets with a probability and p-value above a customizable threshold.

In total, we obtained thousands of predictions, the majority of which have low probabilities (≤0.5) (See Additional Datafiles 3-7). To focus our attention on likely sRNA-target pairs without excluding too many predictions we settled on targets with a predicted probability ≥0.7, referred to as probable going forward. We also define the term MPT, most probable target, as the prediction given the highest probability across all predictions for a given sRNA. Lastly, we define a consensus target as a gene that was predicted to be the MPT for an sRNA in at least four of the six *S. pneumoniae* strains. This term only pertains to the predictions made by IntaRNA, sRNARFTarget, and TargetRNA3. We observed none of the sRNARFTarget predictions are probable. This in combination with our baseline evaluation of the csRNAs led us to focus on the predictions made by IntaRNA, TargetRNA3, and CopraRNA when applicable.

In many cases sRNA pairing can impact the internal base pairing of the mRNA to enable gene expression changes (Papenfort and Storz 2024). Thus, we assessed whether the putative mRNA target regions are likely to have internal base pairing that may be impacted sRNA interaction. We used RNAfold (Lorenz et al., 2011) to predict the structure of mRNA target sequences including 25 nucleotides up/downstream of the binding region in the absence of any sRNA partner. We consider a structured region to be a segment of the mRNA sequence displaying internal base pairing (>2 consecutive bases). Across the MPTs predicted by IntaRNA and TargetRNA3, 51 out of 58 sRNAs base pair with a region considered structured suggesting that the putative sRNA-mRNA interaction may induce structural changes in the secondary or tertiary structures to enable regulation (Tieng et al., 2023).

### sRNAs may play a role in pathogenesis via their targets

Previous studies used transposon insertion mutants to conclude specific sRNAs may support virulence in TIGR4 (Mann et al., 2012). We compared our predicted targets for these sRNAs to evaluate these hypotheses. Previous work suggested 8 putative sRNAs (Table 2) play a definitive role in pathogenesis, and some individual target loci were identified by microarray analysis of attenuated sRNA mutants (Mann et al., 2012). Only 3 of these sRNAs met our criteria for further investigation (Fig. 2A, Table 2). The others overlap with known *cis*-regulatory elements (F20 and F44), transfer-messenger RNA (F32), or were removed following conservation analysis (F41, F48). F41 is one of the sRNAs found in <12 strains, and F48 was deemed a repetitive sequence (average of 36 copies per genome). Among the three remaining candidates, one (M1/F7) is a csRNA with an established role in pathogenesis (DeLay 2024, Patenge et al., 2012, Schnorpfeil et al., 2013). The other two, M45 and M23, are not characterized. M45 is predicted to target type IV teichoic acid flippase TacF that is responsible for transporting choline across the cytoplasmic membrane, a nutritional requirement of *S. pneumoniae* (Damjanovic et al., 2007). M23 was originally reported to target SP_RS08340-50, a putative carbohydrate transporter, based on microarray analysis, but we predict that it targets a transposase encoding transcript (SP_RS13320) (Table 2). Without validating the targets, the role of M45 and M23 in pathogenesis is unclear, however the predicted target (*tacF*) of M45 is suggestive of such a role.

**Table 2:**
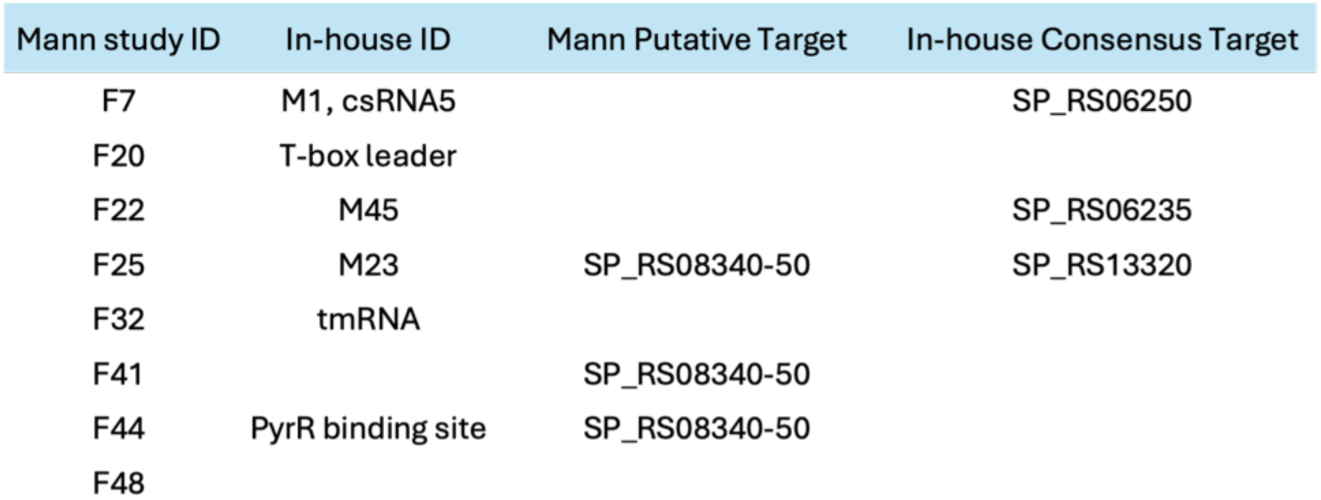
A comparison of the in-house target predictions and the putative targets identified by microarray analysis. The previously identified putative targets SP_RS08340-50 are three neighboring loci involved in carbohydrate transport (carbohydrate ABC transporter permease) and proposed to be collectively regulated by three of the sRNAs (Mann et al., 2012).

To further assess whether specific sRNAs are potentially regulating multiple targets in a previously recognized regulatory response (Table 3), we investigated whether the predicted targets belong to established operons or regulons in TIGR4. The regulons that appeared the most often are PyrR, CodY, and CcpA and we noticed the targets belonging to established operons are always the first or last gene in the operon with the first gene being more common. This suggests that the sRNAs may be inhibiting translation, typically blocking the ribosome binding site (e.g. start of an operon), or stabilizing the transcript by binding to the 3’ end of the mRNA, depending on the relative location of interaction (Papenfort and Vanderpool, 2015). Most notably one sRNA, M63 (Fig. 3A), is predicted to target four different genes in the CcpA regulon (Table 3), genes regulated by the catabolite control protein A (CcpA), an essential transcription factor in Gram-positive bacteria that is responsible for mediating carbon catabolite repression and activation. In *S. pneumoniae*, mutations in CcpA reduce virulence in mouse models (Giammarinaro & Paton, 2002; Iyer et al., 2005). We also see that M63 interacts with the different targets in various regions of the sRNA with unique base-pairing (Fig. 3B). Lastly, M63 is unique in that it is the only sRNA candidate where every reported target has a probability ≥0.7. Furthermore, this is true for all 6 *S. pneumoniae* strains assessed. M63 also overlaps with the rio16 sORF, indicating that part of this sRNA is translated. The sORF has a conserved sequence with average sequence identity of 99.7% across the pangenome. However, it remains possible that M63 is a dual-function RNA both encoding a small protein and regulating the members of the CcpA regulon. The observation that M63 has multiple probable targets acting on a regulon associated with virulence (Iyer et al. 2005) makes this sRNA a high priority candidate for further investigation.

**Figure 3:**
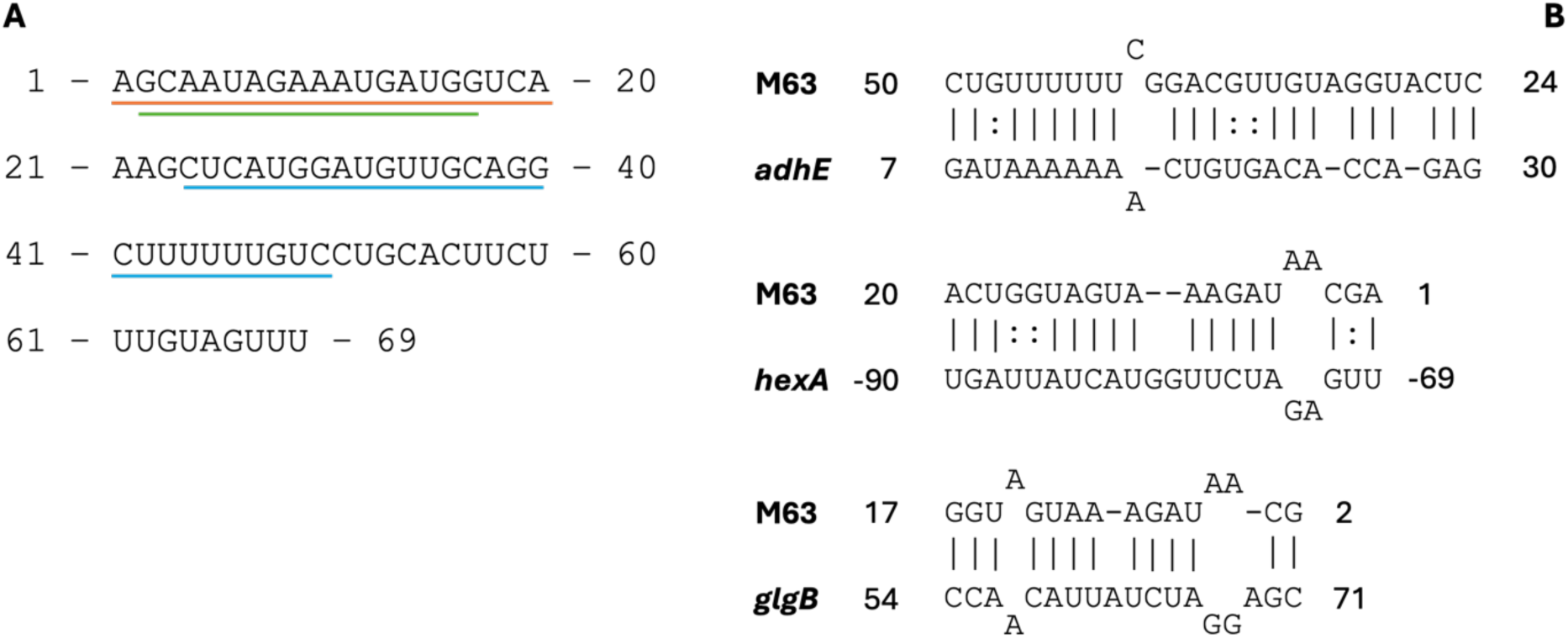
**A)** The M63 sequence with underlined subsequences indicating the regions interacting in part B. Blue corresponds to the *adhE* interaction, orange to *hexA*, and green to *glgB.* **B)** Three of the M63 interactions predicted by TargetRNA3 in TIGR4. The gene names are predicted targets of M63 in the CcpA regulon. The numbers on either end are relative positions of the interacting sequences within the full RNA sequence for the sRNA. For the mRNAs, numbering is relative to the mRNA start codon. A “:” indicates a G-U pair.

**Table 3:**
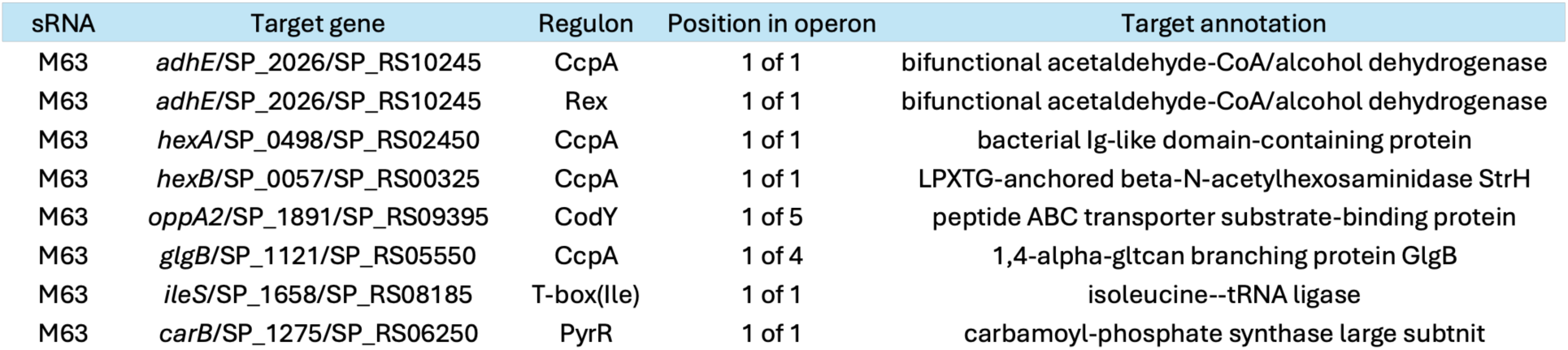
The regulons, according to the RegPrecise database, in which all targets of M63 are involved. Predictions made by TargetRNA3 in *S. pneumoniae* strain TIGR4.

### Transposase associated sRNAs are frequent

Among the TargetRNA3 predictions, we noticed a large number of transposase associated targets. Candidates targeting transposases encoding transcripts include M10, M47, M62, and M69. These sRNAs are all encoded antisense to an annotated transposase (IL3 or IL30 family), overlapping with the 5’-UTR or first few amino acids of the gene. A subset of these, M10, M47 and M62, show substantial sequence identity to each other, with M10 having a 3’-extention compared to M47 and M62 (Fig. 4A). Notably, M10, M47, and M62 all have a large number of BLAST hits in the genome, but these sequences did not exceed our threshold of >15 hits in the genome to be considered repetitive sequences. M47 and M62 have consensus targets with both IntaRNA and TargetRNA3 predicting transposase encoding targets. The collected targets for this set of sRNAs (M10, M47, and M62) includes over 18 different transposase genes, with all but two interactions showing high probability ≥0.92 (Supplemental Table S2). This large number of targets results from the duplicated nature of the sequence immediately surrounding the transposase gene (Fig. 4B). However, we note that all but one of the targets of M62 have a predicted probability below our threshold of 0.7, potentially the result of the differences in length between the M62 sequence and the M10 and M47 sequences (region accessibility due to RNA folding is a factor in target prediction and may be influenced by the extra flanking sequence).

**Figure 4:**
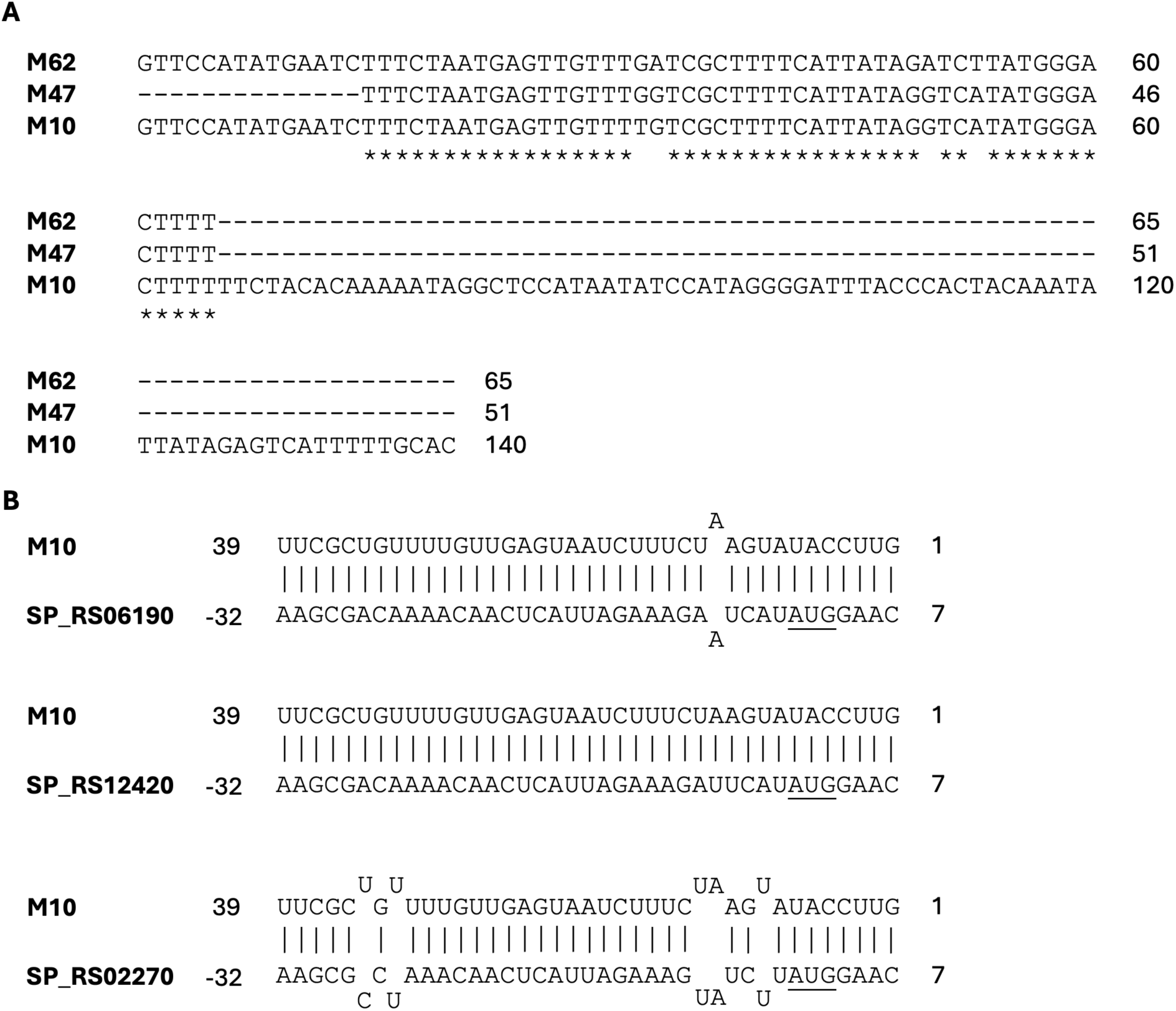
**A)** Clustal omega alignments of M10, M47, and M62 which are predicted to target numerous transposase encoding genes. ‘*’ indicates a position in which all three sequences overlap. **B)** Three of the M10 interactions predicted by TargetRNA3 in TIGR4. The numbers on either end are relative positions of the interacting sequences within the full RNA sequence. A negative number indicates the sequence is upstream of the mRNA start codon. SP_RS### is the TIGR4 target locus. The underlined “AUG”s are the transposase gene start codons.

There are several well-characterized examples of transposon antisense encoded RNAs in bacteria including RNA-OUT, inhibiting IS10 (Simons and Kleckner, 1983), art200, inhibiting IS200 (Ellis et al., 2015), and RNA-C inhibiting IS30 (Arini et al., 1997). Transposon associated antisense RNAs that overlap the transposon coding sequence proximal to the start codon typically act in trans, blocking the translation of the transposase (Simons and Kleckner 1983, Ellis, 2015), but RNA-C, which is antisense to the transposase gene but not directly at the start codon, only appears to act in cis (Arini, 1997). Thus, based on the position of these sRNAs proximal and overlapping the ribosome-binding site, it is likely that they are *trans*-acting across the many transposon copies present in the genome. The sequence of M69 is distinct from that of M10, M47, and M62; however, its placement upstream and antisense to an annotated transposase gene suggests a similar functionality.

### *Cis*-encoded sRNA candidates are less common

To identify other *cis-*encoded sRNA candidates we compared the genomic coordinates of the candidates and MPTs. Candidates suspected to be *cis*-encoded must have genomic coordinates overlapping with the target coordinates. IntaRNA shows 19 candidates may be *cis*-encoded, three of which exhibit probable interaction (Fig. 5A). In contrast, TargetRNA3 suggests only a single candidate to be *cis*-encoded, and it is probable and in common with IntaRNA’s results (Fig. 5B). The possible *cis*-encoded sRNA predicted by TargetRNA3 is M66 (Fig. 5C). The M66 sequence appears antisense to the target with perfect binding across 40 nucleotides. M66 targets *ruvB* that codes for the Holliday junction branch migration DNA helicase RuvB, a subunit in the RuvABC complex. The complex processes Holliday junctions, nucleic acid structures that contain four joined double-stranded arms, during genetic recombination and DNA repair. The individual RuvB subunit is a hexameric ring helicase that acts like a motor to draw the DNA through the complex (Sharples et al., 1999). We believe that the overlap in *cis*-encoded predictions made by TargetRNA3 and IntaRNA suggest that this target is one of interest.

**Figure 5:**
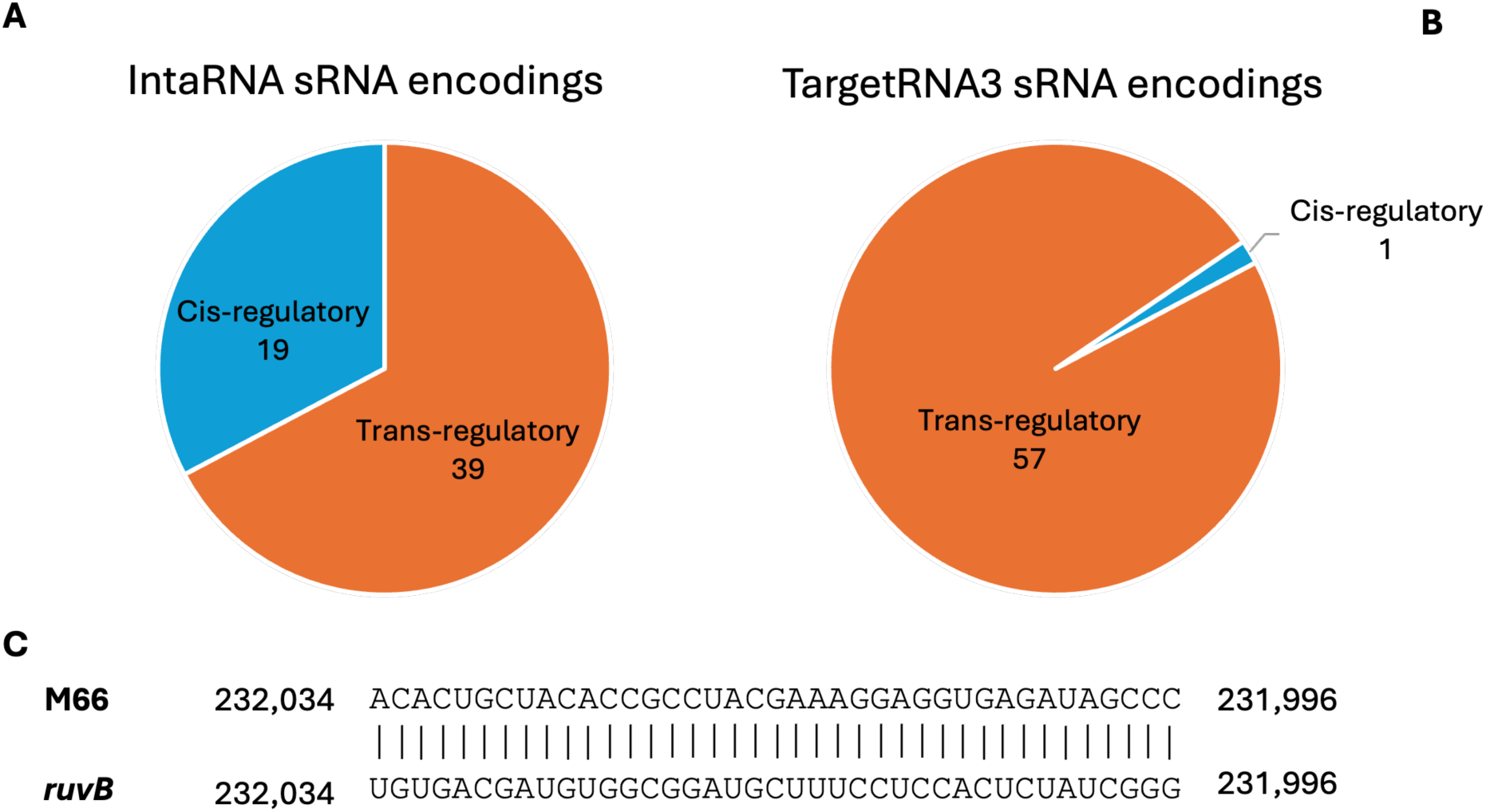
**A)** The distribution of sRNA locations relative to its targets predicted by IntaRNA for 58 candidate sRNAs **B)** The distribution of sRNA locations relative to its targets predicted by TargetRNA3. **C)** The M66 cis-interaction is predicted by TargetRNA3 and the sRNA is found on the complement strand and the target on the top strand. The numbers on either end are exact positions of the interacting sequences in the D39 genome (NC_008533.2).

### Seven notable sRNAs for future experimental validation

Across the 58 sRNAs, 7 stood out for reasons that we believe warrant future work to experimentally validate this study’s results. Each sRNA is highly conserved and targets a gene with high probability. 4 of the notable sRNA candidates share a consensus target between at least two RIP programs (M18, M47, M62, M66) (Table 4). We believe a consensus target, a gene predicted to be the MPT for an sRNA in at least four of the six *S. pneumoniae* strains, is indicative of a highly likely true sRNA-target pair. Four of the notable candidates target multiple transposases (M10, M47, M62, M69). In addition to transposon associated sRNAs, there are also sRNA candidates with potential metabolic targets such as M63, which has 13 probable targets four of which are in the CcpA regulon. We also checked PneumoBrowse2 (Janssen et al., 2024), an interactive online platform with detailed annotations of *S. pneumoniae* genomes like D39V, D39W and TIGR4, for additional annotations on our 7 notable sRNAs. M18 is annotated as part of an anti-toxin/toxin system (Type I addiction module toxin, Fst family) and our predicted target is the ClC family H(+)/Cl(-) exchange transporter. From our analysis, we speculate that these 7 sRNAs are the most likely to lead to future validation of true sRNA-target pairs that may inform us about the role of sRNAs in *S. pneumoniae* metabolism and virulence.

**Table 4:** Notable sRNAs with a high degree of conservation and interesting predicted targets. Other IDs are the labels assigned in the data source studies. The “Consensus” column refers to whether a consensus target was found across multiple RIP programs for the sRNA. The “Mechanism” column refers to whether the sRNA is suspected to be *cis*-acting or *trans*-acting.

| In-house sRNA ID | Other IDs | Consensus | Predicted target(s) | Mechanism | Additional Comments |
| --- | --- | --- | --- | --- | --- |
| M63 | Spd_sr37/srf-04 |  | CcpA regulon | trans | Possibly influences virulence via CcpA regulon |
| M10 | srn266/Spd_sr55 |  | transposases | trans | Similar targets to M47, many transposon related |
| M47 | F31 | ✓ | transposases | trans | Similar targets to M10, many transposon related |
| M62 | Spd_sr6/srf-03 | ✓ | transposases | trans | RNA sequence related to M10 and M47 |
| M69 | Spd_sr47 |  | transposases | trans | Targets 2 transposases with high probability |
| M66 | Spd_sr24 | ✓ | *ruvB* | cis | *ruvB* involved in recombination and DNA repair |
| M18 | srn503/F67/Spd_sr111 | ✓ | SP_RS05710 | trans | Targets an H(+)/Cl(-) exchange transporter |

## Conclusion

This study compiled a list of sRNAs in *S. pneumoniae*, then analyzed their conservation across the Streptococcus genus and within the *S. pneumoniae* pangenome and predicted mRNA targets of conserved sRNAs. BLAST indicates 58 sRNAs exhibit strong conservation across 385 strains. Four RNA-RNA interaction prediction programs made thousands of predictions for the 58 sRNAs. Ultimately, only the probable targets predicted by IntaRNA and TargetRNA3 were the focus of this study’s target examination. From our analysis we propose that there are a handful of transposon associated sRNAs that target transposases encoding transcripts, likely acting in trans based on the position of the base-pairing. However, we also identified high probability targets for other sRNA candidates. For example, M63 (Spd_sr37/srf-04), has several predicted targets across the CcpA regulon implying a potential role in carbon metabolism. Through this work, we have identified a list of 7 sRNAs for which biological function can be hypothesized. Future work will strive to experimentally validate these hypotheses to reveal more regarding the nature of these sRNAs, their targets in *S. pneumoniae*, and roles in growth and virulence.

## Methods

### Compiling sRNA data sources

Previous studies identified putative sRNAs by high-throughput sequencing in *S. pneumoniae* strains TIGR4 (NC_003028.3) and D39W (CP000410.1). The putative sRNAs in TIGR4 (Acebo et al., 2012, Mann et al., 2012) and D39W (Sinha et al., 2019) were narrowed down to a new list of candidates for further analysis of conservation and target prediction (see Results and Discussion).

### Creating the in-house list of putative sRNAs

A list of in-house putative sRNAs was created from the candidates. Previous TIGR4 studies identified different coordinates for transcription initiation and termination sites, so new in-house coordinates were created by combining the smallest initiation and largest termination site coordinates (Supplemental Fig. S1A). In-house Python scripts retrieved the sRNA sequences from the TIGR4 and D39W genomes using the new coordinates. sRNAs identified in multiple studies under different names were assigned an in-house ID and sequence using the new coordinates (e.g. csRNA5/SN35/srn061/F7/CcnE becomes M1). Differences between the TIGR4 and D39W genomes forced the need to compare sequences rather than coordinates. VectorBuilder (https://en.vectorbuilder.com/tool/sequence-alignment.html) aligned sRNAs to confirm the sequences overlap. The largest possible sequence became the new in-house sRNA by joining the overlapping subsequence and trailing sequences on either end (Supplemental Fig. S1B). After forming the list of in-house sRNAs, sequences with length <50 nucleotides were removed.

### Assessing conservation of sRNAs

The in-house sRNAs were aligned to the genomes of 385 *S. pneumoniae* strains (Cremers et al., 2015, Rosconi et al., 2022), *S. pyogenes* (NZ_LS483338.1), *S. mutans* (NZ_CP044221.1), *S. suis* (NC_012926.1), *S. mitis* (NZ_GL397179.1), *S. oralis* (NZ_LR134336.1), and *S. gordonii* (NZ_CP077224.1). Raw reads and the mapping results for 350 *S. pneumoniae* strains (Cremers et al., 2015) available in BAM format were converted to consensus files in FASTA format with the samtools consensus mode (Danecek et al., 2021). Then, a database, created with makeblastdb, containing the sRNA sequences was aligned to each genome using blastn with the task parameter set to megablast. Similarly, a database created with makeblastdb containing the 385 *S. pneumoniae* strains was aligned to each of the sORF nucleotide and peptide sequences using blastn and tblastn. The e-value maximum threshold was raised from a default of 10 to 20 for the tblastn alignments. An in-house Python script retrieved the average number of alignments, best sequence identity, and number of genomes with an alignment for each sRNA. In-house sRNAs with an average number of alignments per strain >15 were classified as potentially highly repetitive sequences and removed from the in-house list. sRNAs not appearing in the majority of the 385 strains were also removed. A database containing the *S. pyogenes*, *S. mutans*, *S. suis*, *S. mitis*, *S. oralis*, and *S. gordonii* genomes, created with makeblastdb, was used to search for sRNA sequence alignments using blastn with the task parameter set to blastn. Synteny of the sRNAs was evaluated by comparing the 1000 nt upstream and downstream of the sequences to those in TIGR4. The 1000 nt on either end of the sRNA sequences for the other 384 strains were compared base-wise to obtain sequence identities. The average sequence identities, both up and downstream, were averaged across all 384 strains, and if either average sequence identity, up or downstream, is ≥75% then it was concluded that the synteny of the sRNA is preserved.

### sRNA target prediction

IntaRNA version 3.3.2, sRNARFTarget, and TargetRNA3 were used to predict targets for the 58 sRNA candidates. CopraRNA version 2.1.4 was only used to predict targets for 3 sRNAs with significant alignments in *S. suis*, *S. pyogenes*, *S. mutans, S. mitis, S. oralis,* and *S. gordonii*. Predictions for each sRNA were made in 6 different *S. pneumoniae* strains consisting of TIGR4 (NC_003028.3), D39 (NC_008533.2), TVO_Taiwan19F-14, TVO_1901920, TVO_1901934, TVO_1902277 (Rosconi et al., 2022). IntaRNA was run with the IntaRNAsTar personality, the number of predictions set to 5, and otherwise default parameters. sRNARFTarget was run with the provided Docker container. sRNARFTarget requires a user input transcriptome, so an in-house Python script created two transcriptome files for each of the six *S. pneumoniae* strains using the coordinates from the respective GenBank files and retrieving the sequences from the genome. The two transcriptomes are defined by the exact gene coordinates and coordinates adjusted to include 100 nucleotides upstream of the start codon and 300 nucleotides downstream of the stop codon or until the next gene’s start codon, whichever comes first. TargetRNA3 was run with the probability threshold lowered to 0.25 and otherwise default parameters. Note, the six genomes were first added to the local user database by providing the accession identifier. CopraRNA was run with default parameters in *S. pneumoniae* (NC_003028), *S. suis* (NC_012926), *S. pyogenes* (NZ_LS483338), *S. mutans* (NZ_CP044221), *S. mitis* (NZ_CP012646), *S. oralis* (NZ_LR134336), and *S. gordonii* (NZ_CP077224).

### sRNA target analysis

An in-house Python script retrieved the sequences including 25 nucleotides upstream and downstream of the mRNA interacting sequence. The structures of these sequences were predicted using RNAfold version 2.6.4 with default parameters. If the sRNA interacting sequence overlaps with a structured region of the mRNA, then the interaction was labeled as interacting with a structured region. Only the MPTs of each sRNA in the six strains predicted by IntaRNA were analyzed. To determine if the sRNAs are acting in regulons we searched the target loci against the RegPrecise database (https://regprecise.lbl.gov/index.jsp). Only the MPTs and probable targets predicted by TargetRNA3 in TIGR4 were searched seeing as TIGR4 is the only *S. pneumoniae* strain for which the database contains information. The expression of sRNA sequences was confirmed by checking if the transcription initiation and termination sites were present in 5’ and 3’ RNA-end sequencing data (Warrier et al., 2018).

## Supporting information

SupplementalFile2_python_scripts

Additional Datafile 3

Additional Datafile 1

Additional Datafile 7

Additional Datafile 2

Additional Datafile 6

Additional Datafile 5

Additional Datafile 4

## Additional Data Files

Additional_Data_1_source-sRNAs.xlsx – initial list of putative sRNAs from three studies

Additional_Data_2_In-house-sRNAs.xlsx – final in-house list after deduplication (Fig. 1C)

Additional_Data_3_CopraRNA.xlsx – CopraRNA target predictions

Additional_Data_4_IntaRNA.xlsx – IntaRNA target predictions

Additional_Data_5_sRNARFTarget-Transcriptome-Exact.xlsx – sRNARFT target predictions (coding sequences only)

Additional_Data_6_sRNARFTarget-Transcriptome-100-300.xlsx – sRNARFT target predictions (coding sequences and flanking regions)

Additional_Data_7_TargetRNA3.xlsx – TargetRNA3 target predictions

## Supplemental Files

Eichelman_SupplementalFile1.pdf – Supplemental Figures and Tables

Eichelman_SupplementalFile2_python_scripts.zip – In-house python scripts

## Acknowledgments

MCE was partially funded by a Boston College Undergraduate Research Fellowship and by grants R21AI181123 and R01GM134259 from the US National Institutes of Health. We would like to thank Quinlan Furumo for careful copyediting of the manuscript.

MCE roles: data curation, investigation, visualization, writing – original draft

MMM roles: conceptualization, data curation, funding acquisition, supervision, writing – review and editing

## References

Acebo, Paloma, Antonio J. Martin-Galiano, Sara Navarro, Ángel Zaballos, and Mónica Amblar. 2012. “Identification of 88 Regulatory Small RNAs in the TIGR4 Strain of the Human Pathogen *Streptococcus Pneumoniae*.” RNA 18 (3): 530–46. 10.1261/rna.027359.111.

Altschul, Stephen F., Warren Gish, Webb Miller, Eugene W. Myers, and David J. Lipman. 1990. “Basic Local Alignment Search Tool.” Journal of Molecular Biology 215 (3): 403–10. 10.1016/S0022-2836(05)80360-2.

Aoyama, Jordan J., and Gisela Storz. 2023. “Two for One: Regulatory RNAs That Encode Small Proteins.” Trends in Biochemical Sciences 48 (12): 1035–43. 10.1016/j.tibs.2023.09.002.

Arini, Achille, Marcel P. Keller, and Werner Arber. 1997. “An Antisense RNA in IS30 Regulates the Translational Expression of the Transposase.” Biological Chemistry 378 (12). 10.1515/bchm.1997.378.12.1421.

Babina, Arianne M., Mark W. Soo, Yang Fu, and Michelle M. Meyer. 2015. “An S6:S18 Complex Inhibits Translation of *E. Coli rpsF*.” RNA 21 (12): 2039–46. 10.1261/rna.049544.115.

Busch, Anke, Andreas S. Richter, and Rolf Backofen. 2008. “IntaRNA: Efficient Prediction of Bacterial sRNA Targets Incorporating Target Site Accessibility and Seed Regions.” Bioinformatics 24 (24): 2849–56. 10.1093/bioinformatics/btn544.

CDC, National Center for Emerging Zoonotic and Infectious Diseases (U.S.), National Center for HIV/AIDS, Viral Hepatitis, STD, and TB Prevention (U.S.), and National Center for Immunization and Respiratory Diseases (U.S.). 2013. Antibiotic Resistance Threats in the United States, 2013. https://stacks.cdc.gov/view/cdc/20705/cdc_20705_DS1.pdf.

Christopoulou, Niki, and Sander Granneman. 2022. “The Role of RNA-Binding Proteins in Mediating Adaptive Responses in Gram-Positive Bacteria.” The FEBS Journal 289 (7): 1746–64. 10.1111/febs.15810.

Cremers, Amelieke J. H., Fredrick M. Mobegi, Marien I. De Jonge, Sacha A. F. T. Van Hijum, Jacques F. Meis, Peter W. M. Hermans, Gerben Ferwerda, Stephen D. Bentley, and Aldert L. Zomer. 2015. “The Post-Vaccine Microevolution of Invasive Streptococcus Pneumoniae.” Scientific Reports 5 (1): 14952. 10.1038/srep14952.

Damjanovic, Marlen, Arun S. Kharat, Alice Eberhardt, Alexander Tomasz, and Waldemar Vollmer. 2007. “The Essential *tacF* Gene Is Responsible for the Choline-Dependent Growth Phenotype of *Streptococcus Pneumoniae*.” Journal of Bacteriology 189 (19): 7105–11. 10.1128/JB.00681-07.

Danecek, Petr, James K Bonfield, Jennifer Liddle, John Marshall, Valeriu Ohan, Martin O Pollard, Andrew Whitwham, et al. 2021. “Twelve Years of SAMtools and BCFtools.” GigaScience 10 (2): giab008. 10.1093/gigascience/giab008.

De Lay, Nicholas R., Nidhi Verma, Dhriti Sinha, Abigail Garrett, Maximillian K. Osterberg, Daisy Porter, Spencer Reiling, David P. Giedroc, and Malcolm E. Winkler. 2024. “The Five Homologous CiaR-Controlled Ccn sRNAs of Streptococcus Pneumoniae Modulate Zn-Resistance.” Edited by Rachel M. McLoughlin. PLOS Pathogens 20 (10): e1012165. 10.1371/journal.ppat.1012165.

Ellis, Michael J., Ryan S. Trussler, and David B. Haniford. 2015. “A *Cis* -Encoded sRNA, Hfq and mRNA Secondary Structure Act Independently to Suppress IS *200* Transposition.” Nucleic Acids Research 43 (13): 6511–27. 10.1093/nar/gkv584.

Felden, Brice, and Yoann Augagneur. 2021. “Diversity and Versatility in Small RNA-Mediated Regulation in Bacterial Pathogens.” Frontiers in Microbiology 12 (August):719977. 10.3389/fmicb.2021.719977.

Fu, Yang, Kaila Deiorio-Haggar, Jon Anthony, and Michelle M. Meyer. 2013. “Most RNAs Regulating Ribosomal Protein Biosynthesis in Escherichia Coli Are Narrowly Distributed to Gammaproteobacteria.” Nucleic Acids Research 41 (6): 3491–3503. 10.1093/nar/gkt055.

Furumo, Quinlan, and Michelle M. Meyer. 2024. “PIPETS: A Statistically Informed, Gene-Annotation Agnostic Analysis Method to Study Bacterial Termination Using 3′-End Sequencing.” BMC Bioinformatics 25 (1): 363. 10.1186/s12859-024-05982-5.

GBD 2016 Lower Respiratory Infections Collaborators. 2018. “Estimates of the Global, Regional, and National Morbidity, Mortality, and Aetiologies of Lower Respiratory Infections in 195 Countries, 1990-2016: A Systematic Analysis for the Global Burden of Disease Study 2016.” The Lancet. Infectious Diseases 18 (11): 1191–1210. 10.1016/S1473-3099(18)30310-4.

Giammarinaro, Philippe, and James C. Paton. 2002. “Role of RegM, a Homologue of the Catabolite Repressor Protein CcpA, in the Virulence of *Streptococcus Pneumoniae*.” Infection and Immunity 70 (10): 5454–61. 10.1128/IAI.70.10.5454-5461.2002.

Halfmann, Alexander, Márta Kovács, Regine Hakenbeck, and Reinhold Brückner. 2007. “Identification of the Genes Directly Controlled by the Response Regulator CiaR in *Streptococcus Pneumoniae* : Five out of 15 Promoters Drive Expression of Small Non-coding RNAs.” Molecular Microbiology 66 (1): 110–26. 10.1111/j.1365-2958.2007.05900.x.

Hör, Jens, Geneviève Garriss, Silvia Di Giorgio, Lisa-Marie Hack, Jens T. Vanselow, Konrad U. Förstner, Andreas Schlosser, Birgitta Henriques-Normark, and Jörg Vogel. 2020. “Grad-Seq in a Gram-Positive Bacterium Reveals Exonucleolytic sRNA Activation in Competence Control.” The EMBO Journal 39 (9): e103852. 10.15252/embj.2019103852.

Huang, Susan S., Kristen M. Johnson, G. Thomas Ray, Peter Wroe, Tracy A. Lieu, Matthew R. Moore, Elizabeth R. Zell, et al. 2011. “Healthcare Utilization and Cost of Pneumococcal Disease in the United States.” Vaccine 29 (18): 3398–3412. 10.1016/j.vaccine.2011.02.088.

Iyer, Ramkumar, Nitin S. Baliga, and Andrew Camilli. 2005. “Catabolite Control Protein A (CcpA) Contributes to Virulence and Regulation of Sugar Metabolism in *Streptococcus Pneumoniae*.” Journal of Bacteriology 187 (24): 8340–49. 10.1128/JB.187.24.8340-8349.2005.

Jabbour, Nancy, and Marie-Frédérique Lartigue. 2021. “An Inventory of CiaR-Dependent Small Regulatory RNAs in Streptococci.” Frontiers in Microbiology 12 (May):669396. 10.3389/fmicb.2021.669396.

Janssen, Axel B, Paddy S Gibson, Afonso M Bravo, Vincent de Bakker, Jelle Slager, and Jan-Willem Veening. 2024. “PneumoBrowse 2: An Integrated Visual Platform for Curated Genome Annotation and Multiomics Data Analysis of *Streptococcus Pneumoniae*.” Nucleic Acids Research, October, gkae923. 10.1093/nar/gkae923.

Laczkovich, Irina, Kyle Mangano, Xinhao Shao, Adam J. Hockenberry, Yu Gao, Alexander Mankin, Nora Vázquez-Laslop, and Michael J. Federle. 2022. “Discovery of Unannotated Small Open Reading Frames in Streptococcus Pneumoniae D39 Involved in Quorum Sensing and Virulence Using Ribosome Profiling.” Edited by N. Luisa Hiller. mBio 13 (4): e01247–22. 10.1128/mbio.01247-22.

Li, Wuju, Xiaomin Ying, Qixuan Lu, and Linxi Chen. 2012. “Predicting sRNAs and Their Targets in Bacteria.” Genomics, Proteomics & Bioinformatics 10 (5): 276–84. 10.1016/j.gpb.2012.09.004.

Lorenz, Ronny, Stephan H Bernhart, Christian Höner Zu Siederdissen, Hakim Tafer, Christoph Flamm, Peter F Stadler, and Ivo L Hofacker. 2011. “ViennaRNA Package 2.0.” Algorithms for Molecular Biology 6 (1): 26. 10.1186/1748-7188-6-26.

Mann, Beth, Tim Van Opijnen, Jianmin Wang, Caroline Obert, Yong-Dong Wang, Robert Carter, Daniel J. McGoldrick, et al. 2012. “Control of Virulence by Small RNAs in Streptococcus Pneumoniae.” Edited by Pascale Cossart. PLoS Pathogens 8 (7): e1002788. 10.1371/journal.ppat.1002788.

Mann, Martin, Patrick R. Wright, and Rolf Backofen. 2017. “IntaRNA 2.0: Enhanced and Customizable Prediction of RNA–RNA Interactions.” Nucleic Acids Research 45 (W1): W435–39. 10.1093/nar/gkx279.

Martin, Julia E., My T. Le, Nabin Bhattarai, Daiana A. Capdevila, Jiangchuan Shen, Malcolm E. Winkler, and David P. Giedroc. 2019. “A Mn-Sensing Riboswitch Activates Expression of a Mn2+/Ca2+ ATPase Transporter in Streptococcus.” Nucleic Acids Research 47 (13): 6885–99. 10.1093/nar/gkz494.

Marx, Patrick, Michael Nuhn, Martá Kovács, Regine Hakenbeck, and Reinhold Brückner. 2010. “Identification of Genes for Small Non-Coding RNAs That Belong to the Regulon of the Two-Component Regulatory System CiaRH in Streptococcus.” BMC Genomics 11 (1): 661. 10.1186/1471-2164-11-661.

Mascher, Thorsten, Manuel Heintz, Dorothea Zähner, Michelle Merai, and Regine Hakenbeck. 2006. “The CiaRH System of *Streptococcus Pneumoniae* Prevents Lysis during Stress Induced by Treatment with Cell Wall Inhibitors and by Mutations in *Pbp2x* Involved in β-Lactam Resistance.” Journal of Bacteriology 188 (5): 1959–68. 10.1128/JB.188.5.1959-1968.2006.

Naskulwar, Kratika, and Lourdes Peña-Castillo. 2022. “sRNARFTarget: A Fast Machine-Learning-Based Approach for Transcriptome-Wide sRNA Target Prediction.” RNA Biology 19 (1): 44–54. 10.1080/15476286.2021.2012058.

Olejniczak, Mikolaj, Xiaofang Jiang, Maciej M. Basczok, and Gisela Storz. 2022. “KH Domain Proteins: Another Family of Bacterial RNA Matchmakers?” Molecular Microbiology 117 (1): 10–19. 10.1111/mmi.14842.

Papenfort, Kai, and Gisela Storz. 2024. “Insights into Bacterial Metabolism from Small RNAs.” Cell Chemical Biology 31 (9): 1571–77. 10.1016/j.chembiol.2024.07.002.

Papenfort, Kai, and Carin K. Vanderpool. 2015. “Target Activation by Regulatory RNAs in Bacteria.” FEMS Microbiology Reviews 39 (3): 362–78. 10.1093/femsre/fuv016.

Patenge, Nadja, Tomas Fiedler, and Bernd Kreikemeyer. 2012. “Common Regulators of Virulence in Streptococci.” In Host-Pathogen Interactions in Streptococcal Diseases, edited by G. Singh Chhatwal, 368:111–53. Current Topics in Microbiology and Immunology. Berlin, Heidelberg: Springer Berlin Heidelberg. 10.1007/82_2012_295.

Peer, Asaf, and Hanah Margalit. 2014. “Evolutionary Patterns of *Escherichia Coli* Small RNAs and Their Regulatory Interactions.” RNA 20 (7): 994–1003. 10.1261/rna.043133.113.

Raden, Martin, Syed M Ali, Omer S Alkhnbashi, Anke Busch, Fabrizio Costa, Jason A Davis, Florian Eggenhofer, et al. 2018. “Freiburg RNA Tools: A Central Online Resource for RNA-Focused Research and Teaching.” Nucleic Acids Research 46 (W1): W25–29. 10.1093/nar/gky329.

Raina, Medha, Alisa King, Colleen Bianco, and Carin K. Vanderpool. 2018. “Dual-Function RNAs.” Microbiology Spectrum 6 (5). 10.1128/microbiolspec.RWR-0032-2018.

Rosconi, Federico, Emily Rudmann, Jien Li, Defne Surujon, Jon Anthony, Matthew Frank, Dakota S. Jones, et al. 2022. “A Bacterial Pan-Genome Makes Gene Essentiality Strain-Dependent and Evolvable.” Nature Microbiology 7 (10): 1580–92. 10.1038/s41564-022-01208-7.

Schnorpfeil, Anke, Miriam Kranz, Martá Kovács, Christian Kirsch, Julia Gartmann, Ines Brunner, Sabrina Bittmann, and Reinhold Brückner. 2013. “Target Evaluation of the Non-coding CSRNAS Reveals a Link of the Two-component Regulatory System CIARH to Competence Control in *S Treptococcus Pneumoniae* R 6.” Molecular Microbiology 89 (2): 334–49. 10.1111/mmi.12277.

Sharples, Gary J., Stuart M. Ingleston, and Robert G. Lloyd. 1999. “Holliday Junction Processing in Bacteria: Insights from the Evolutionary Conservation of RuvABC, RecG, and RusA.” Journal of Bacteriology 181 (18): 5543–50. 10.1128/JB.181.18.5543-5550.1999.

Simons, Robert W., and Nancy Kleckner. 1983. “Translational Control of IS10 Transposition.” Cell 34 (2): 683–91. 10.1016/0092-8674(83)90401-4.

Sinha, Dhriti, Kurt Zimmer, Todd A. Cameron, Douglas B. Rusch, Malcolm E. Winkler, and Nicholas R. De Lay. 2019. “Redefining the Small Regulatory RNA Transcriptome in Streptococcus Pneumoniae Serotype 2 Strain D39.” Edited by Victor J. DiRita. Journal of Bacteriology 201 (14). 10.1128/JB.00764-18.

Tieng, Francis Yew Fu, Muhammad-Redha Abdullah-Zawawi, Nur Alyaa Afifah Md Shahri, Zeti-Azura Mohamed-Hussein, Learn-Han Lee, and Nurul-Syakima Ab Mutalib. 2023. “A Hitchhiker’s Guide to RNA–RNA Structure and Interaction Prediction Tools.” Briefings in Bioinformatics 25 (1): bbad421. 10.1093/bib/bbad421.

Tjaden, Brian. 2023. “TargetRNA3: Predicting Prokaryotic RNA Regulatory Targets with Machine Learning.” Genome Biology 24 (1): 276. 10.1186/s13059-023-03117-2.

Tsui, Ho-Ching Tiffany, Dhriti Mukherjee, Valerie A. Ray, Lok-To Sham, Andrew L. Feig, and Malcolm E. Winkler. 2010. “Identification and Characterization of Noncoding Small RNAs in *Streptococcus Pneumoniae* Serotype 2 Strain D39.” Journal of Bacteriology 192 (1): 264–79. 10.1128/JB.01204-09.

Warrier, Indu, Nikhil Ram-Mohan, Zeyu Zhu, Ariana Hazery, Haley Echlin, Jason Rosch, Michelle M. Meyer, and Tim Van Opijnen. 2018. “The Transcriptional Landscape of Streptococcus Pneumoniae TIGR4 Reveals a Complex Operon Architecture and Abundant Riboregulation Critical for Growth and Virulence.” Edited by Carlos Javier Orihuela. PLOS Pathogens 14 (12): e1007461. 10.1371/journal.ppat.1007461.

Wright, Patrick R., Jens Georg, Martin Mann, Dragos A. Sorescu, Andreas S. Richter, Steffen Lott, Robert Kleinkauf, Wolfgang R. Hess, and Rolf Backofen. 2014. “CopraRNA and IntaRNA: Predicting Small RNA Targets, Networks and Interaction Domains.” Nucleic Acids Research 42 (W1): W119–23. 10.1093/nar/gku359.

Wright, Patrick R., Andreas S. Richter, Kai Papenfort, Martin Mann, Jörg Vogel, Wolfgang R. Hess, Rolf Backofen, and Jens Georg. 2013. “Comparative Genomics Boosts Target Prediction for Bacterial Small RNAs.” Proceedings of the National Academy of Sciences 110 (37). 10.1073/pnas.1303248110.

Zhang, Aixia, Karen M. Wassarman, Carsten Rosenow, Brian C. Tjaden, Gisela Storz, and Susan Gottesman. 2003. “Global Analysis of Small RNA and mRNA Targets of Hfq.” Molecular Microbiology 50 (4): 1111–24. 10.1046/j.1365-2958.2003.03734.x.

Zorgani, Mohamed A., Roland Quentin, and Marie-Frédérique Lartigue. 2016. “Regulatory RNAs in the Less Studied Streptococcal Species: From Nomenclature to Identification.” Frontiers in Microbiology 7 (July). 10.3389/fmicb.2016.01161.

