## SupplementalFIle1_Supplemental_Figures&Tables for "Assessing the conservation and targets of putative sRNAs in *Streptococcus pneumoniae*"

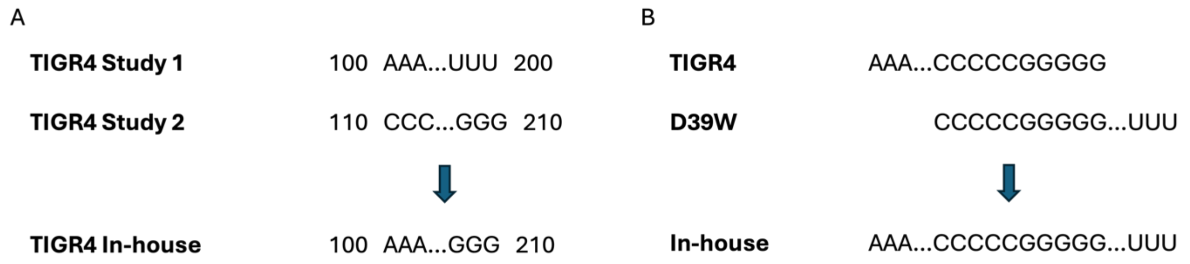


**Figure S1**: **A)** Formation of the in-house TIGR4 sRNA sequences by combining the sequences identified in the two TIGR4 studies. **B)** Formation of the in-house sRNA sequences by combining the sequences identified in TIGR4 and D39W studies.

| | sRNA conservation in other Streptococcus species | | | | | |
|---|---|---|---|---|---|---|
| | S. mitis | S. oralis | S. gordonii | S. pyogenes | S. mutans | S. suis |
| M1 | | | | | | |
| M4 | | | | | | |
| M5 | | | | | | |
| M6 | | | | | | |
| M7 | X | X | | | | |
| M9 | X | X | | | | |
| M10 | | | | X | X | X |
| M13 | | | | | | |
| M14 | X | X | | | | |
| M16 | | | | | | |
| M17 | | | | | | |
| M18 | | | | | | |
| M19 | X | X | X | | | |
| M20 | X | X | | | | |
| M22 | | | | | | |
| M23 | | | | | | |
| M24 | | | | | | |
| M25 | | | | | | |
| M29 | X | X | | | | |
| M30 | X | | | | X | |
| M32 | | | | | | |
| M34 | | | | | | |
| M36 | | X | | | | |
| M37 | | | | | | |
| M39 | | | | | | X |
| M40 | | X | | | | |
| M41 | | | | | | |
| M42 | | | | | | |
| M43 | X | | | | | |
| M44 | | | | | | X |
| M45 | X | | | | | |
| M46 | | | | | | |
| M47 | | | | X | X | X |
| M48 | | | | | | |
| M49 | | | | | | |
| M50 | | | | | | |
| M52 | X | X | | | | |
| M53 | | X | | | | |
| M59 | | | | | | |
| M61 | | | | | | |
| M62 | | | | X | X | X |
| M63 | X | X | X | X | | |
| M64 | | | | | | |
| M65 | X | X | | | | |
| M66 | | | | | | |
| M67 | X | X | | | | |
| M68 | X | | | | | |
| M69 | | | | X | | |
| M70 | | X | | | | |
| M71 | | | | | | |
| M73 | | | | | | |
| M74 | | | | | | |
| M75 | | | | | | |
| M76 | X | X | | | | |
| M77 | X | X | X | X | X | X |
| M79 | | | | | | |
| M80 | | | | | | |
| M81 | X | X | X | X | X | X |

**Table S1**: The conservation of the 58 sRNA candidates in 6 other Streptococcus species. “X” indicates the sRNA in conserved in that species.

| sRNA | Target locus | Annotation | Probability | Interaction Length (nt) |
|---|---|---|---|---|
| M10 | SP_RS06190 | ISL3 family transposase | 0.960 | 40 |
| M10 | SP_RS08360 | ISL3 family transposase | 0.953 | 40 |
| M10 | SP_RS12115 | ISL3 family transposase | 0.952 | 40 |
| M10 | SP_RS11735 | ISL3 family transposase | 0.952 | 40 |
| M10 | SP_RS12420 | transposase | 0.949 | 40 |
| M10 | SP_RS12405 | ISL3 family transposase | 0.949 | 40 |
| M10 | SP_RS12565 | transposase family protein | 0.946 | 40 |
| M10 | SP_RS11780 | ISL3 family transposase | 0.944 | 40 |
| M10 | SP_RS04090 | ISL3-like element IS1167A family transposase | 0.941 | 40 |
| M10 | SP_RS05040 | ISL3 family transposase | 0.941 | 40 |
| M10 | SP_RS11845 | transposase | 0.934 | 40 |
| M10 | SP_RS00665 | ISL3 family transposase | 0.930 | 40 |
| M10 | SP_RS02270 | ISL3 family transposase | 0.929 | 40 |
| M10 | SP_RS11995 | ISL3 family transposase | 0.926 | 40 |
| M10 | SP_RS08895 | ISL3 family transposase | 0.925 | 40 |
| M10 | SP_RS08080 | ISL3 family transposase | 0.924 | 40 |
| M10 | SP_RS07795 | ISL3-like element IS1167A family transposase | 0.922 | 40 |
| M10 | SP_RS07095 | ISL3 family transposase | 0.922 | 40 |
| M47 | SP_RS06190 | ISL3 family transposase | 0.960 | 39 |
| M47 | SP_RS11735 | ISL3 family transposase | 0.952 | 39 |
| M47 | SP_RS12115 | ISL3 family transposase | 0.952 | 39 |
| M47 | SP_RS08360 | ISL3 family transposase | 0.950 | 39 |
| M47 | SP_RS12405 | ISL3 family transposase | 0.949 | 39 |
| M47 | SP_RS12420 | transposase | 0.949 | 39 |
| M47 | SP_RS12565 | transposase family protein | 0.946 | 39 |
| M47 | SP_RS11780 | ISL3 family transposase | 0.944 | 39 |
| M47 | SP_RS04090 | ISL3-like element IS1167A family | 0.941 | 39 |
| M47 | SP_RS05040 | ISL3 family transposase | 0.941 | 39 |
| M47 | SP_RS11845 | transposase | 0.938 | 38 |
| M47 | SP_RS11995 | ISL3 family transposase | 0.926 | 39 |
| M47 | SP_RS07095 | ISL3 family transposase | 0.922 | 39 |
| M47 | SP_RS08080 | ISL3 family transposase | 0.921 | 39 |
| M47 | SP_RS08895 | ISL3 family transposase | 0.919 | 39 |
| M47 | SP_RS07795 | ISL3-like element IS1167A family | 0.916 | 39 |
| M47 | SP_RS00665 | ISL3 family transposase | 0.722 | 39 |
| M47 | SP_RS02270 | ISL3 family transposase | 0.710 | 39 |
| M62 | SP_RS11845 | transposase | 0.734 | 40 |

**Table S2**: The M10, M47, and M62 target predictions in TIGR4 predicted by TargetRNA3. Only the predictions with a probability over 0.7 are shown for each sRNA. M62 only has one predicted transposase target with a probability over 0.7.
