## SupplementalFile2_python_scripts for "Assessing the conservation and targets of putative sRNAs in *Streptococcus pneumoniae*": check_rna_orf.py

```
"""
Get the sequences of the sRNA
including 10 nucleotides upstream.
These sequences will be used to
check if a start codon is present
immediately upstream.
"""


import os
from typing import IO
import helper_functions as hf


def write_to_outfile(outfile: IO[str], data: tuple[str, str]) -> None:
    """Write the data to the output file"""
    name: str = data[0]
    sequence: str = data[1]
    outfile.write(f'>{name}\n{sequence}\n\n')


def parse_coords(line: str) -> tuple[str, list[str], str]:
    """Parse the coordinates into a start and end"""
    data: list[str] = []
    name: str = ''
    tigr4: str = ''
    d39: str = ''
    if len(line.split('\t')) < 4:
        data = line.split('\t')
        name = data[0]
        tigr4 = data[2]
    else:
        data = line.split('\t')
        name = data[0]
        tigr4 = data[2]
        d39 = data[3]
    if len(tigr4) < 1:
        return (name, d39.strip().split('-'), 'd39')
    return (name, tigr4.split('-'), 'tigr4')


def main() -> None:
    """The main function"""
    path: str = os.getcwd()[:-7]
    coords_file: str = os.path.join(path, 'sRNA data/sRNA coords.txt')
    d39_fa: str = os.path.join(path, 'Other fasta files/D39.fasta')
    tigr4_fa: str = os.path.join(path, 'Other fasta files/TIGR4.fasta')
    srnas_fa: str = os.path.join(path, 'sRNA data/In-house_sRNAsV4.fasta')
    ten_nt_up_file: str = os.path.join(path, 'sRNA data/sRNA-58-10ntUp.fa')
    with (open(coords_file, 'r', encoding='utf-8') as infile1,
          open(tigr4_fa, 'r', encoding='utf-8') as infile2,
          open(d39_fa, 'r', encoding='utf-8') as infile3,
          open(srnas_fa, 'r', encoding='utf-8') as infile4,
          open(ten_nt_up_file, 'w', encoding='utf-8') as outfile):

        for line in infile1:
            srna = infile4.readline()
            while line.split('\t')[0] not in srna:
                srna = infile4.readline()
            seq = infile4.readline().strip()
            strand = srna.strip()[-1]
            name: str = ''
            coords: list[str] = []
            strain: str = ''
            name, coords, strain = parse_coords(line)
            start: int = 0
            end: int = 0
            start, end = int(coords[0]), int(coords[1])
            sequence = ''
            if strand == '+':
                if strain == 'd39':
                    sequence = hf.retrieve_sequence(start, start, infile3, 10 - 1, 0)
                else:
                    sequence = hf.retrieve_sequence(start, start, infile2, 10 - 1, 0)
            else:
                if strain == 'd39':
                    sequence = hf.retrieve_sequence(end, end, infile3, 0, 10 - 1)
                else:
                    sequence = hf.retrieve_sequence(end, end, infile2, 0, 10 - 1)
                sequence = hf.reverse_complement(sequence)
            write_to_outfile(outfile, (name, sequence + seq))


if __name__ == '__main__':
    main()
```
