## SupplementalFile2_python_scripts for "Assessing the conservation and targets of putative sRNAs in *Streptococcus pneumoniae*": coprarna_top_hits.py

```
"""
Organize the top 5 hits of each sRNA for
CopraRNA into a single .csv that can
be easily copy/pasted into excel
"""


import os
from typing import IO


def write_data(information: list[str], outfile: IO[str]) -> None:
    """Write the data to the output file"""
    information[5] = information[5][:-1].split('|') # type: ignore
    for item in information:
        if item is information[5]: # type: ignore
            if len(item) == 1:
                for _ in range(8):
                    outfile.write('N/A;')
            else:
                for info in information[5]:
                    outfile.write(str(info) + ';')
        else:
            outfile.write(str(item) + ';')
    outfile.write('\n')
    # sRNA_ID
    # Rank
    # Organism
    # CopraRNA p-val
    # Locus tag
    # Gene name
    # Energy
    # IntaRNA p-val
    # mRNA_start
    # mRNA_end
    # sRNA_start
    # sRNA_end
    # Gene_ID
    # Annotation


def main() -> None:
    """The main function"""
    path: str = os.getcwd()[:-7]
    srna_id: str = 'M77'
    directory: str = path + 'CopraRNA stuff/New-Iteration/' + srna_id + '/'
    infilename: str = directory + srna_id + '_result.csv'
    outfilename: str = directory + srna_id + '_summary.csv'

    organisms: dict[int, str] = {2: 'S. pneumoniae (TIGR4)',
                       3: 'S. mutans',
                       4: 'S. suis',
                       5: 'S. pyogenes',
                       6: 'S. oralis',
                       7: 'S. gordonii'}
    # Change depending on the number of input species

    # data = [fdr,
    #         p-value,            CopraRNA p-value
    #         NC_003028,          pneumoniae (TIGR4)
    #         NZ_CP044221,        mutans
    #         NC_012926,          suis
    #         NZ_LS483338,        pyogenes
    #         NZ_LR134336         oralis
    #         NZ_CP077224         gordonii
    #         Annotation,
    #         Additional homologs,
    #         Amount sampled
    #         ]

    with (open(infilename, 'r', encoding='utf-8') as infile,
          open(outfilename, 'w', encoding='utf-8') as outfile):
        infile.readline()
        header = ('sRNA_ID;Rank;Organism;CopraRNA_p-val;Locus_tag;'
                  'Gene_name;Energy;IntaRNA_p-val;mRNA_start;mRNA_end;'
                  'sRNA_start;sRNA_end;Gene_ID;Annotation\n')
        outfile.write(header)
        for rank in range(1, 6):
            data: list[str] = infile.readline().split(',')
            for i in range(2, 8):
                locus: str = ''
                organism: str = ''
                try:
                    locus, organism = data[i].split('(')
                except ValueError:
                    locus, organism = 'N/A', 'N/A'
                n: int = list(organisms.keys())[-1] + 1
                information: list[str] = [srna_id, str(rank), organisms[i],
                                          data[1], locus, organism, data[n]]
                write_data(information, outfile)


if __name__ == '__main__':
    main()
```
