## SupplementalFile2_python_scripts for "Assessing the conservation and targets of putative sRNAs in *Streptococcus pneumoniae*": evaluate_synteny.py

```
"""
Evaluate the synteny of the sRNAs by comparing
the sequences 1000 nt up and downstream of the
sRNA sequences for each of the 385 strains
"""


import os
import helper_functions as hf


def compare_seqs(seq1: str, seq2: str) -> float:
    """Compare two sequences and return the percentage similarity"""
    mismatch: int = 0
    for i in range(min(len(seq1), len(seq2))):
        if seq1[i] != seq2[i]:
            mismatch += 1
    return round((1 - mismatch / len(seq1)) * 100, 2)


def get_coords(filename: str, srna: str) -> tuple[str, str]:
    """Get the coordinates of the sRNA in the genome"""
    with open(filename, 'r', encoding='utf-8') as infile:
        # Confirm there is a hit for the sRNA first
        hit: bool = False
        line: str = infile.readline()
        query: str = ''
        last_query: str = ''
        temp: str = ''
        start: str = ''
        end: str = ''
        while line[0] != '>':
            if srna + ':' in line:
                hit = True
            line = infile.readline()
        infile.seek(infile.tell() - 100)
        if not hit:
            return ('', '')
        line = infile.readline()
        while line.split(':')[0] != ('>' + srna):
            line = infile.readline()
        for _ in range(6):
            infile.readline()
        _, start, _, temp = infile.readline().split('  ')
        while True:
            for _ in range(3):
                infile.readline()
            query = infile.readline()
            if 'Query' not in query:
                break
            last_query = query
        try:
            end = last_query.strip().split('  ')[-1] or temp
        except ValueError:
            end = temp
    return (start, end)


def main() -> None:
    """Main function"""
    path: str = os.getcwd()[:-7]
    fastas: list[str] = []
    for filename in os.listdir(path + 'Other fasta files'):
        if filename[-6:] == '.fasta' and 'ignore' not in filename:
            fastas.append(filename)
    for filename in os.listdir(path + 'TVO fasta files'):
        if filename[-6:] == '.fasta':
            fastas.append(filename)
    for filename in os.listdir(path + 'Clinical fasta files'):
        if filename[-6:] == '.fasta':
            fastas.append(filename)

    srnas: list[str] = []
    line: str = ''
    with open(path + 'sRNA data/In-house_sRNAsV3.fasta', 'r', encoding='utf-8') as infile:
        line = infile.readline()
        while line:
            srnas.append(line[1:].split(':')[0])
            for _ in range(2):
                infile.readline()
            line = infile.readline()

    with (open(path + 'sRNA data/synteny_results.txt', 'w', encoding='utf-8') as outfile,
          open(path + 'Other fasta files/TIGR4.fasta', 'r', encoding='utf-8') as infile):
        outfile.write('sRNA, Is the sequence preserved (Yes/No)')
        for srna in srnas:
            avg_up: float = 0
            avg_down: float = 0
            strains: int = 0
            data: tuple[str, str] = get_coords(path + 'sRNA BLAST search results/TIGR4results.txt',
                                               srna)
            start: str = data[0]
            end: str = data[1]
            tigr4_up: str = hf.retrieve_sequence(int(start) - 1000, int(start) - 1, infile)
            tigr4_down: str = hf.retrieve_sequence(int(end) + 1, int(end) + 1000, infile)
            for strain in fastas:
                file: str = path + 'sRNA BLAST search results/' + strain[:-6] + 'results.txt'
                start, end = get_coords(file, srna)
                if start == '' and end == '':
                    continue
                strains += 1
                up: str = ''
                down: str = ''
                if '#' in strain:
                    with (open(path + 'Clinical fasta files/' + strain, 'r', encoding='utf-8')
                          as infile):
                        up = hf.retrieve_sequence(int(start) - 1000, int(start) - 1, infile)
                        down = hf.retrieve_sequence(int(end) + 1, int(end) + 1000, infile)
                elif 'TVO' in strain:
                    with (open(path + 'TVO fasta files/' + strain, 'r', encoding='utf-8')
                          as infile):
                        up = hf.retrieve_sequence(int(start) - 1000, int(start) - 1, infile)
                        down = hf.retrieve_sequence(int(end) + 1, int(end) + 1000, infile)
                else:
                    with (open(path + 'Other fasta files/' + strain, 'r', encoding='utf-8')
                          as infile):
                        up = hf.retrieve_sequence(int(start) - 1000, int(start) - 1, infile)
                        down = hf.retrieve_sequence(int(end) + 1, int(end) + 1000, infile)
                avg_up += compare_seqs(tigr4_up, up)
                avg_down += compare_seqs(tigr4_down, down)
            avg_up /= strains
            avg_down /= strains
            outfile.write(srna)
            if avg_up >= 75:
                outfile.write(', Yes')
            else:
                outfile.write(', No')
            if avg_down >= 75:
                outfile.write(', Yes\n')
            else:
                outfile.write(', No\n')


if __name__ == '__main__':
    main()
```
