## SupplementalFile2_python_scripts for "Assessing the conservation and targets of putative sRNAs in *Streptococcus pneumoniae*": get_d39_seqs.py

```
"""
Get the sequences for the putative sRNAs from the
D39W study (Sinha) using the provided coordinates
This list is only contains the putative sRNAs that
were also found in the other studies (at least 1
other study) or confirmed by Northern/qRT-PCR
D39W genome: CP000410.1
"""


import os
import helper_functions as hf


def main() -> None:
    """The main function"""
    path: str = os.getcwd()[:-7]
    d39: str = path + 'Other fasta files/D39W.fasta'
    d39_srnas: str = path + 'sRNA data/D39W_sRNAs.txt'
    outfilename: str = path + 'sRNA data/D39W_seqs.fasta'
    with (open(d39, 'r', encoding='utf-8') as infile1,
          open(d39_srnas, 'r', encoding='utf-8') as infile2,
          open(outfilename, 'w', encoding='utf-8') as outfile):

        hf.skip_lines(infile2, 2)
        line: str = infile2.readline()
        while True:
            if not line:
                break
            d39_name: str = ''
            study_names: str = ''
            coordinates: str = ''
            strand: str = ''
            d39_name, study_names, coordinates, strand = line.split(' ')
            strand = strand[1]
            start, end = coordinates.split('-')
            sequence: str = hf.retrieve_sequence(int(start), int(end), infile1)
            if strand == '-':
                sequence = hf.reverse_complement(sequence)
            outfile.write(f'>{d39_name}|{study_names}|{start}-{end}|{strand}\n{sequence}\n\n')
            line = infile2.readline()


if __name__ == '__main__':
    main()
```
