## SupplementalFile2_python_scripts for "Assessing the conservation and targets of putative sRNAs in *Streptococcus pneumoniae*": get_mann_seqs.py

```
"""
Get the sequences for the putative sRNAs from the
Mann study (TIGR4) using the provided coordinates
TIGR4 genome: NC_003028.3
"""


import os
import helper_functions as hf


def main() -> None:
    """The main function"""
    path: str = os.getcwd()[:-7]
    tigr4: str = path + 'Other fasta files/TIGR4.fasta'
    mann_coords: str = path + 'sRNA data/Mann_coords.txt'
    outfilename: str = path + 'sRNA data/Mann_seqs.fasta'
    with (open(tigr4, 'r', encoding='utf-8') as infile1,
          open(mann_coords, 'r', encoding='utf-8') as infile2,
          open(outfilename, 'w', encoding='utf-8') as outfile):

        hf.skip_lines(infile2, 2)
        line: str = infile2.readline()
        while line:
            if not line:
                break
            name: str = ''
            coordinates: str = ''
            name, coordinates = line.split('\t')
            strand: str = name[0]
            start, end = coordinates.split('-')
            sequence: str = hf.retrieve_sequence(int(start), int(end), infile1)
            if strand == 'R':
                sequence = hf.reverse_complement(sequence)
            outfile.write(f'>{name}|{start}-{end}{sequence}\n\n')
            line = infile2.readline()


if __name__ == '__main__':
    main()
```
