## SupplementalFile2_python_scripts for "Assessing the conservation and targets of putative sRNAs in *Streptococcus pneumoniae*": get_old_loci.py

```
"""
Find the old loci names of the TargetRNA3
TIGR4 targets for analyzing the distribution
of location
"""


import os
from typing import IO


def get_locus(infile: IO[str], locus: str) -> str:
    """Get the locus name from the GenBank file"""
    infile.seek(0)
    while locus not in infile.readline():
        continue
    try:
        tag = infile.readline().split('"')[1]
    except IndexError:
        tag = ''
    return tag


def main() -> None:
    """The main function"""
    path: str = os.getcwd()[:-7]
    targetrna: str = path + 'TargetRNA3 stuff/'
    strain: str = 'TIGR4'
    txt1: str = targetrna  + 'All_TIGR4_new_loci.txt'
    txt2: str = targetrna  + 'All_TIGR4_new_loci_highprob.txt'
    genbank: str = path + 'Annotated genomes/' + strain + '.gb'
    # These correspond to the regular file
    output1: str = targetrna + '/All_TIGR4_old_loci_withdupes1.txt'
    output2: str = targetrna + '/All_TIGR4_old_loci_nodupes1.txt'
    # These correspond to the high prob file
    output3: str = targetrna + '/All_TIGR4_old_loci_withdupes2.txt'
    output4: str = targetrna + '/All_TIGR4_old_loci_nodupes2.txt'

    new_loci_nd1: list[str] = []
    new_loci_wd1: set[str] = set()
    locus: str = ''
    old_locus: str = ''
    with open(txt1, 'r', encoding='utf-8') as infile:
        locus = infile.readline().strip()
        while locus:
            if ' ' in locus:
                locus = locus.split(' ')[1][1:-1]
            new_loci_nd1.append(locus)
            new_loci_wd1.add(locus)
            locus = infile.readline().strip()

    new_loci_nd2: list[str] = []
    new_loci_wd2: set[str] = set()
    with open(txt2, 'r', encoding='utf-8') as infile:
        locus = infile.readline().strip()
        while locus:
            if ' ' in locus:
                locus = locus.split(' ')[1][1:-1]
            new_loci_nd2.append(locus)
            new_loci_wd2.add(locus)
            locus = infile.readline().strip()

    with open(genbank, 'r', encoding='utf-8') as infile:
        with open(output1, 'w', encoding='utf-8') as outfile:
            for locus in new_loci_nd1:
                old_locus = get_locus(infile, locus)
                if old_locus != '':
                    outfile.write(old_locus + '\n')
        with open(output2, 'w', encoding='utf-8') as outfile:
            for locus in new_loci_wd1:
                old_locus = get_locus(infile, locus)
                if old_locus != '':
                    outfile.write(old_locus + '\n')
        with open(output3, 'w', encoding='utf-8') as outfile:
            for locus in new_loci_nd2:
                old_locus = get_locus(infile, locus)
                if old_locus != '':
                    outfile.write(old_locus + '\n')
        with open(output4, 'w', encoding='utf-8') as outfile:
            for locus in new_loci_wd2:
                old_locus = get_locus(infile, locus)
                if old_locus != '':
                    outfile.write(old_locus + '\n')


if __name__ == '__main__':
    main()
```
