## SupplementalFile2_python_scripts for "Assessing the conservation and targets of putative sRNAs in *Streptococcus pneumoniae*": get_tigr4_seqs.py

```
"""
Get the sequences for the putative sRNAs from the
TIGR4 studies (Mann and Acebo) using the provided
coordinates
This list is only contains the putative sRNAs that
were also found in the other studies (at least 1
other study) or confirmed by Northern/qRT-PCR
TIGR4 genome: NC_003028.3
"""


import os
import helper_functions as hf


def main() -> None:
    """The main function"""
    path: str = os.getcwd()[:-7]
    tigr4: str = path + 'Other fasta files/TIGR4.fasta'
    tigr4_srnas: str = path + 'sRNA data/TIGR4_sRNAs.txt'
    outfilename: str = path + 'sRNA data/TIGR4_seqs.fasta'
    with (open(tigr4, 'r', encoding='utf-8') as infile1,
          open(tigr4_srnas, 'r', encoding='utf-8') as infile2,
          open(outfilename, 'w', encoding='utf-8') as outfile):

        hf.skip_lines(infile2, 2)
        line: str = infile2.readline()
        while True:
            if not line:
                break
            tigr4_name: str = ''
            study_names: str = ''
            coordinates: str = ''
            strand: str = ''
            start: str = ''
            end: str = ''
            tigr4_name, study_names, coordinates, strand = line.split(' ')
            strand = strand[1]
            start, end = coordinates.split('-')
            sequence: str = hf.retrieve_sequence(int(start), int(end), infile1)
            if strand == '-':
                sequence = hf.reverse_complement(sequence)
            outfile.write(f'>{tigr4_name}|{study_names}|{start}-{end}|{strand}\n{sequence}\n\n')
            line = infile2.readline()


if __name__ == '__main__':
    main()
```
