## SupplementalFile2_python_scripts for "Assessing the conservation and targets of putative sRNAs in *Streptococcus pneumoniae*": helper_functions.py

```
"""
A file that contains all of the functions
used in the scripts
"""


from typing import IO


def complement(seq: str) -> str:
    """Return the complement sequence of the input sequence"""
    seq = seq.replace('A', '?')
    seq = seq.replace('T', 'A')
    seq = seq.replace('?', 'T')
    seq = seq.replace('C', '?')
    seq = seq.replace('G', 'C')
    seq = seq.replace('?', 'G')
    return seq


def reverse(seq: str) -> str:
    """Return the input sequence reversed/backwards"""
    return seq[::-1]


def reverse_complement(seq: str) -> str:
    """Return the reverse complement of the input sequence"""
    return reverse(complement(seq))


def rna_to_dna(seq: str) -> str:
    """Return the DNA version of the input RNA sequence"""
    return seq.replace('U', 'T')


def dna_to_rna(seq: str) -> str:
    """Return the RNA version of the DNA sequence"""
    return seq.replace('T', 'U')


def retrieve_sequence(start: int, end: int, infile: IO[str], upstream: int = 0,
                      downstream: int = 0) -> str:
    """Return the DNA sequence for given genomic coordinates"""
    seq: str = ''
    infile.seek(0)
    infile.readline()
    line_length: int = len(infile.readline().strip())
    infile.seek(0)
    infile.readline()
    position: int = 1
    num_lines: int = (start - upstream) // line_length
    if start - upstream > 0:
        for _ in range(num_lines):
            infile.readline()
        position = num_lines * line_length + 1
    while position <= start - upstream - 1:
        if infile.read(1) != '\n':
            position += 1
    while position <= end + downstream:
        char: str = infile.read(1)
        if not char:
            break
        if char != '\n':
            seq += char
            position += 1
    return seq


def skip_lines(infile: IO[str], num: int = 1) -> None:
    """Skip some number of lines in a file being read"""
    for _ in range(num):
        infile.readline()
```
