## SupplementalFile2_python_scripts for "Assessing the conservation and targets of putative sRNAs in *Streptococcus pneumoniae*": individual_srna_fastas.py

```
"""
Separate the In-house_sRNAsV3.fasta file
into individual files containing a single
sRNA for using with TargetRNA3
"""


import os


def main() -> None:
    """The main function"""
    path: str = os.getcwd()[:-7]
    individuals: str = path + 'sRNA data/Individual sRNA fastas/'
    srnas: str = path + 'sRNA data/In-house_sRNAsV3.fasta'
    with open(srnas, 'r', encoding='utf-8') as infile:
        line: str = infile.readline()
        while line:
            name: str = line
            sequence: str = infile.readline()
            infile.readline()
            m_name: str = name[1:].split(':')[0]
            outfilename: str = individuals + m_name + '.fasta'
            with open(outfilename, 'w', encoding='utf-8') as outfile:
                outfile.write(f'{name}{sequence}')
            line = infile.readline()


if __name__ == '__main__':
    main()
```
