## SupplementalFile2_python_scripts for "Assessing the conservation and targets of putative sRNAs in *Streptococcus pneumoniae*": intarna_hits.py

```
"""
Find the locus, gene, upstream, etc. data
using the csv output from IntaRNA.
Output as a .tsv so it can be copy pasted
into excel.
"""


import os
from typing import IO
import helper_functions as hf


def get_interaction(bp_list: str) -> str:
    """Generate a string representing the interaction between the sRNA and mRNA"""
    interaction: str = ''
    new_bp_list: list[str] = bp_list.split(':')
    start: int = int(new_bp_list[0].split(',')[0][1:])
    end: int = int(new_bp_list[-1].split(',')[0][1:])
    num_bases: int = end - start + 1
    base_pairs: list[int] = [0] * num_bases
    position: int = 0
    for _, bp in enumerate(new_bp_list):
        position = int(bp.split(',')[0][1:]) - start
        base_pairs[position] = 1
    for i, num in enumerate(base_pairs):
        if i == len(base_pairs) // 2:
            interaction += '&'
        elif num == 1 and i < len(base_pairs) / 2:
            interaction += '('
        elif num == 1 and i > len(base_pairs) / 2:
            interaction += ')'
        else:
            interaction += '.'
    return interaction


def write_header(outfile: IO[str]) -> None:
    """Write the header for the output file"""
    header: str = ('sep=;\n'
              'sRNA_ID;sRNA_subseq_start;sRNA_subseq_end;'
              'target;target_start;target_end;mRNA_subseq_start;'
              'mRNA_subseq_end;energy;probability;annotation;upstream_gene;'
              'upstream_start;upstream_end;downstream_gene;downstream_start;'
              'downstream_end;interaction\n')
    outfile.write(header)


def write_data(data: tuple[str, ...], outfile: IO[str]) -> None:
    """Write the data for the output file"""
    for i, item in enumerate(data):
        if i != len(data) - 1:
            outfile.write(item + ';')
        else:
            outfile.write(item + '\n')


def get_upstream(infile: IO[str]) -> tuple[str, ...]:
    """Get the upstream gene"""
    upstream: str = ''
    line: str = infile.readline()
    while line[5:9] != 'gene':
        line = infile.readline()
    gene_start: str = ''
    gene_end: str = ''
    gene_start, gene_end = line[21:].split('..')
    if 'comp' in gene_start:
        gene_start = gene_start.split('(')[1]
        gene_end = gene_end.split(')')[0]
    if '>' in gene_end:
        gene_end = gene_end[1:]
    if '<' in gene_start:
        gene_start = gene_start[1:]
    line = infile.readline()
    while '/product=' not in line:
        line = infile.readline()
    i: int = 31
    while line[i] != '"':
        if line[i] == '\n':
            line = infile.readline()
            i = 21
            upstream += ' '
        upstream += line[i]
        i += 1
    return (upstream, gene_start, gene_end)


def get_downstream(infile: IO[str]) -> tuple[str, ...]:
    """Get the downstream gene"""
    downstream: str = ''
    line: str = infile.readline()
    while line[5:9] != 'gene':
        line = infile.readline()
    gene_start: str = ''
    gene_end: str = ''
    gene_start, gene_end = line[21:].split('..')
    if 'comp' in gene_start:
        gene_start = gene_start.split('(')[1]
        gene_end = gene_end.split(')')[0]
    if '>' in gene_end:
        gene_end = gene_end[1:]
    if '<' in gene_start:
        gene_start = gene_start[1:]
    line = infile.readline()
    while '/product=' not in line:
        line = infile.readline()
    i: int = 31
    while line[i] != '"':
        if line[i] == '\n':
            line = infile.readline()
            i = 21
            downstream += ' '
        downstream += line[i]
        i += 1
    return (downstream, gene_start, gene_end)


def get_target(infile: IO[str], mrna_start: int, mrna_end: int) -> tuple[str, ...]:
    """Get the target gene"""
    annotation: str = ''
    infile.seek(0)
    prev: int = 0
    while 'FEATURES' not in infile.readline():
        continue
    up: tuple[str, ...] = ()
    down: tuple[str, ...] = ()
    while True:
        line: str = infile.readline()
        if line[5:9] == 'gene':
            curr = infile.tell()
            gene_start: str = ''
            gene_end: str = ''
            gene_start, gene_end = line[21:].split('..')
            if 'comp' in gene_start:
                gene_start = gene_start.split('(')[1]
                gene_end = gene_end.split(')')[0]
            if '>' in gene_end:
                gene_end = gene_end[1:]
            if '<' in gene_start:
                gene_start = gene_start[1:]
            if (int(gene_start) <= mrna_end <= int(gene_end) and
                int(gene_start) <= mrna_start <= int(gene_end)):
                line = infile.readline()
                target: str = ''
                while line[21] == '/' and line[22] != 'p':
                    if 'gene' in line:
                        target = line[21:].split('"')[1]
                        break
                    if 'old' in line:
                        target = line[21:].split('"')[1]
                        break
                    target = line[21:].split('"')[1]
                    line = infile.readline()
                line = infile.readline()
                while '/product=' not in line:
                    line = infile.readline()
                i: int = 31
                while line[i] != '"':
                    if line[i] == '\n':
                        line = infile.readline()
                        i = 21
                        annotation += ' '
                    annotation += line[i]
                    i += 1
                infile.seek(prev - 100)
                up = get_upstream(infile)
                infile.seek(curr)
                down = get_downstream(infile)
                return (target, annotation, gene_start, gene_end,
                        up[0], up[1], up[2], down[0], down[1], down[2])
            if mrna_start < int(gene_start):
                target = 'non-coding'
                infile.seek(prev - 100)
                up = get_upstream(infile)
                down = get_downstream(infile)
                return (target, 'N/A', 'N/A', 'N/A',
                        up[0], up[1], up[2], down[0], down[1], down[2])
            prev = curr
        if not line:
            break
    return ('', '', '', '', '', '', '', '', '', '')


def main() -> None:
    """The main function"""
    path: str = os.getcwd()[:-7]
    intarna: str = path + 'IntaRNA stuff/'
    strains: list[str] = ['D39',
               'TIGR4',
               'TVO_D39',
               'TVO_TIGR4',
               'TVO_1901920',
               'TVO_1901934',
               'TVO_1902277',
               'TVO_Taiwan19F-14']
    for strain in strains:
        csv: str = intarna + strain + ' files/' + 'csRNA1-res.csv'
        genbank: str = path + 'Annotated genomes/' + strain + '.gb'
        top_hit: str = intarna + strain + ' files/csRNA1Top_Hit.csv'
        top_5_hits: str = intarna + strain + ' files/csRNA1Top_5_Hits.csv'
        with (open(csv, 'r', encoding='utf-8') as infile1,
              open(genbank, 'r', encoding='utf-8') as infile2,
              open(top_hit, 'w', encoding='utf-8') as outfile1,
              open(top_5_hits, 'w', encoding='utf-8') as outfile2):

            write_header(outfile1)
            write_header(outfile2)
            hf.skip_lines(infile1)
            i: int = 0
            line: str = infile1.readline().strip()
            while line:
                _, id2, start1, end1, start2, end2, _, _, bp_list, energy, prob = line.split(';')
                interaction: str = get_interaction(bp_list)
                target_info: tuple[str, ...] = get_target(infile2, int(start1), int(end1))
                gene_start: str = target_info[2]
                gene_end: str = target_info[3]
                data = (id2[:-1], target_info[0], gene_start, gene_end,
                        start1, end1, start2, end2,
                        energy, prob, target_info[1],
                        target_info[4], target_info[5], target_info[6],
                        target_info[7], target_info[8], target_info[9],
                        interaction)
                write_data(data, outfile2)
                if i % 5 == 0:
                    write_data(data, outfile1)
                i += 1
                line = infile1.readline().strip()


if __name__ == '__main__':
    main()
```
