## SupplementalFile2_python_scripts for "Assessing the conservation and targets of putative sRNAs in *Streptococcus pneumoniae*": mit_ora_gor_blast.py

```
"""
Report the most significant alignment for each sRNA
in each of the 3 Streptococcus species (mitis, 
oralis, gordonii).
Output the identities and gaps of each top alignment
for each sRNA for each species to a file called
blast_results_summary.txt.
"""


import os
from typing import IO
import helper_functions as hf


def get_top_hit(infile: IO[str]) -> str:
    """Get the top hit for a given sRNA."""
    hf.skip_lines(infile, 4)
    identities: str = infile.readline()[1:]
    hf.skip_lines(infile, 2)
    for _ in range(3):
        identities += infile.readline()
    return identities


def main() -> None:
    """The main function"""
    path: str = os.getcwd()[:-7]
    directory: str = path + 'mitis oralis gordonii/'
    file: str = directory + 'blast_results.txt'
    output: str = directory + 'blast_results_summary.txt'
    with (open(file, 'r', encoding='utf-8') as infile,
          open(output, 'w', encoding='utf-8') as outfile):

        hf.skip_lines(infile, 15)
        while True:
            line: str = infile.readline().strip()
            if 'Database' in line:
                break
            name: str = line.split('=')[1]
            hf.skip_lines(infile)
            length: str = infile.readline().strip().split('=')[1]
            hf.skip_lines(infile, 3)
            outfile.write(f'Name: {name}, Length: {length}\n')
            line = infile.readline()
            streps: list[str] = []
            while line[0] == 'N':
                streps.append(line.split(' ')[2])
                line = infile.readline()
            hf.skip_lines(infile)
            for i, name in enumerate(streps):
                while infile.readline()[1:6] != 'Score':
                    continue
                identities: str = infile.readline()[1:]
                outfile.write(f'Strain: {name}, {identities}')
                hf.skip_lines(infile, 2)
                while infile.readline()[0] != '>' and i < len(streps) - 1:
                    continue
            outfile.write('\n')
            while infile.readline()[:9] != 'Effective':
                continue
            hf.skip_lines(infile, 2)


if __name__ == '__main__':
    main()
```
