## SupplementalFile2_python_scripts for "Assessing the conservation and targets of putative sRNAs in *Streptococcus pneumoniae*": mrna_interacting_tr3.py

```
"""
Using the RNAfold results and interaction
in the TargetRNA3 top hit csv file compare the
structure to determine if basepairing is
occurring in structured regions
"""


import os
import helper_functions as hf


def main() -> None:
    """The main function"""
    path: str = os.getcwd()[:-7]
    targetrna3: str = path + 'TargetRNA3 stuff/'
    strains: list[str] = ['D39',
               'TIGR4',
               'TVO_1901920',
               'TVO_1902277',
               'TVO_Taiwan19F-14']
    for strain in strains:
        csv: str = targetrna3 + strain + '_results/' + strain + '_updated.csv'
        txt: str = targetrna3 + strain + '_results/mRNA_target_structure.txt'
        output: str = targetrna3 + strain + '_results/mRNA-sRNA_interaction.txt'
        with (open(csv, 'r', encoding='utf-8') as infile1,
             (open(txt, 'r', encoding='utf-8')) as infile2,
             (open(output, 'w', encoding='utf-8')) as outfile):

            hf.skip_lines(infile1, 2)
            line: str = infile1.readline().strip()
            parentheses: str = '()'
            count: int = 0
            while line:
                interaction: str = line.split(';')[-1]
                interaction = interaction.split('&')[0]
                name: str = infile2.readline()[1:]
                infile2.readline()
                structure: str = infile2.readline()
                blank: str = ' ' * 25
                interaction = blank + interaction + '\n'
                outfile.write(f'{name}{structure}{interaction}')
                for i, bp in enumerate(interaction):
                    if structure[i] in parentheses and bp in parentheses:
                        count += 1
                        outfile.write('Overlap!\n')
                        break
                outfile.write('\n')
                line = infile1.readline().strip()
        message: str = 'mRNA-sRNA interacting subsequences having basepairing in structured regions'
        print(f'{strain}: {count} {message}')


if __name__ == '__main__':
    main()
```
