## SupplementalFile2_python_scripts for "Assessing the conservation and targets of putative sRNAs in *Streptococcus pneumoniae*": mrna_target_25.py

```
"""
Get the sequences of the sRNA-mRNA interaction
including 25 nucleotides upstream and downstream
of the sequence.
These sequences will be used by RNAFold to
determine if the base pairing is occurring in
regions in which the mRNA is folding.
"""


import os
from typing import IO
import helper_functions as hf


def outfile_write(outfile: IO[str], data: tuple[str, str]) -> None:
    """Write the data to the output file."""
    outfile.write(f'>{data[0]}\n{data[1]}\n\n')


def main() -> None:
    """The main function"""
    path: str = os.getcwd()[:-7]
    targetrna3: str = path + 'TargetRNA3 stuff/'
    strains: list[str] = ['D39',
               'TIGR4',
               'TVO_1901920',
               'TVO_1902277',
               'TVO_Taiwan19F-14']
    for strain in strains:
        csv: str = targetrna3 + strain + '_results/' + strain + '_updated.csv'
        directory: str = ''
        if strain in ('D39', 'TIGR4'):
            directory = path + 'Other fasta files/'
        else:
            directory = path + 'TVO fasta files/'
        fasta: str = directory + strain + '.fasta'
        output: str = targetrna3 + strain + '_results/mRNA_target_+25.fasta'
        with (open(csv, 'r', encoding='utf-8') as infile1,
              open(fasta, 'r', encoding='utf-8') as infile2,
              open(output, 'w', encoding='utf-8') as outfile):

            hf.skip_lines(infile1, 2)
            line: str = infile1.readline().strip()
            while line:
                (rna_id, _, _, _, strand, _, start, _, substar2, subend2,
                 _, _, _, _, _, _, _, _, _, _, _) = line.split(';')
                if strand == 'comp':
                    if int(substar2) < 0:
                        substar2 = substar2[1:]
                    else:
                        substar2 = '-' + substar2
                    if int(subend2) < 0:
                        subend2 = subend2[1:]
                    else:
                        subend2 = '-' + subend2
                    sequence = hf.retrieve_sequence(int(start) + int(subend2),
                                                    int(start) + int(substar2), infile2, 25, 25)
                else:
                    sequence = hf.retrieve_sequence(int(start) + int(substar2),
                                                    int(start) + int(subend2), infile2, 25, 25)
                outfile_write(outfile, (rna_id, sequence))
                # for _ in range(5):
                #     line = infile1.readline().strip()
                line = infile1.readline().strip()


if __name__ == '__main__':
    main()
```
