## SupplementalFile2_python_scripts for "Assessing the conservation and targets of putative sRNAs in *Streptococcus pneumoniae*": rio_prot_blast_analysis.py

```
"""
Count the number of genomes with a hit for each rioRNA.
Report the average identity %.
Output the numbers to a file called rioRNA_similarity.tsv.
"""


import os
import helper_functions as hf


def main() -> None:
    """The main function"""
    path: str = os.getcwd()[:-7]
    infilename: str = path + 'rioRNAs/385strepsdb/rioRNAs-prot-results.txt'
    outfilename: str = path + 'rioRNAs/385strepsdb/rioRNA_similarity.tsv'

    riornas: dict[str, list[float]] = {}

    with open(infilename, 'r', encoding='utf-8') as infile:
        end: int = 0
        while True:
            line: str = infile.readline()
            while 'Query=' not in line:
                line = infile.readline()
                if not line:
                    end = 1
                    break
            if end:
                break
            rna_id: str = line.split(' ')[1]
            if rna_id not in riornas:
                riornas[rna_id] = [0.0, 0.0]
                # index 0 = identity %
                # index 1 = # genomes present in
            hf.skip_lines(infile, 3)
            if 'Sequences' not in infile.readline():
                continue
            hf.skip_lines(infile)
            line = infile.readline()
            while line[0] != '\n':
                riornas[rna_id][1] += 1
                line = infile.readline()
            hf.skip_lines(infile)
            line = infile.readline()
            while line[0] == '>':
                release: int = 0
                line = infile.readline()
                if 'Length' not in line:
                    hf.skip_lines(infile)
                hf.skip_lines(infile, 2)
                line = infile.readline()
                identity: str = line[1:].split(' ')[2]
                numerator: str = identity.split('/')[0]
                denominator: str = identity.split('/')[1]
                riornas[rna_id][0] += (float(numerator) / float(denominator)) * 100
                while '>' not in line:
                    line = infile.readline()
                    if 'Lambda' in line:
                        release = 1
                        break
                if release:
                    break

    with open(outfilename, 'w', encoding='utf-8') as outfile:
        outfile.write('ID\tAvg ident %\t# gnms in\n')
        for rna_id, value in riornas.items():
            if value[1] != 0:
                value[0] /= value[1]
            outfile.write(rna_id + '\t')
            outfile.write(str(round(value[0], 2)) + '\t')
            outfile.write(str(value[1]) + '\t')
            outfile.write('\n')


if __name__ == '__main__':
    main()
```
