## SupplementalFile2_python_scripts for "Assessing the conservation and targets of putative sRNAs in *Streptococcus pneumoniae*": srna_blast_analysis.py

```
"""
Count the average number of hits for each sRNA.
A high number of average hits would indicate the
sRNA is potentially a BOX element/transposon.
Also report the average identity %.
Output the numbers to a file called sRNA_similarity.txt.
"""


import os
import helper_functions as hf


def main() -> None:
    """The main function"""
    path: str = os.getcwd()[:-7]
    # blast_results: str = path + 'sRNA BLAST search results'
    blast_results: str = path + 'rioRNAs/BLAST results'
    # outfilename: str = path + 'sRNA data/sRNA_similarity.tsv'
    outfilename: str = path + 'rioRNAs/rioRNA_similarity.tsv'

    srnas: dict[str, list[float]] = {}

    for filename in os.listdir(blast_results):
        if filename[-11:] == 'results.txt':
            file: str = blast_results + '/' + filename
            with open(file, 'r', encoding='utf-8') as infile:
                line: str = infile.readline()
                empty: bool = False
                while 'Sequences' not in line:
                    line = infile.readline()
                    if not line:
                        empty = True
                        break
                if empty:
                    continue
                hf.skip_lines(infile)
                line = infile.readline()
                # while line[0] == 'M':
                while line[0] == 'r':
                    # id = line.split(':')[0]
                    rna_id: str = line.split(' ')[0]
                    if rna_id not in srnas:
                        srnas[rna_id] = [0.0, 0.0, 1.0, 0.0, 0.0]
                        # index 0 = # hits
                        # index 1 = identity %
                        # index 2 = # genomes present in
                        # index 3 = # same strand hits
                        # index 4 = best alignment identity
                    else:
                        srnas[rna_id][2] += 1
                    line = infile.readline()
                hf.skip_lines(infile)
                line = infile.readline()
                while line[0] == '>':
                    save: bool = True
                    # id, ignore, sRNA_strand = line.split(' ')
                    rna_id = line.split(' ')[0]
                    # id = id[1:-1]
                    rna_id = rna_id[1:]
                    # sRNA_strand = sRNA_strand.strip()
                    hf.skip_lines(infile, 2)
                    line = infile.readline()
                    while line[0] == ' ':
                        line = infile.readline()
                        if 'Identities' in line:
                            srnas[rna_id][0] += 1
                        identity: str = line[1:].split(' ')[2]
                        numerator: str = identity.split('/')[0]
                        denominator = identity.split('/')[1]
                        srnas[rna_id][1] += (int(numerator) / int(denominator)) * 100
                        if save:
                            srnas[rna_id][4] += (int(numerator) / int(denominator)) * 100
                            save = False
                        hf.skip_lines(infile)
                        # genome_strand = infile.readline().strip().split('/')[1]
                        # if ((genome_strand == 'Minus' and sRNA_strand == '-')
                        #     or (genome_strand == 'Plus' and sRNA_strand == '+')):
                        #     srnas[id][3] += 1
                        hf.skip_lines(infile)
                        while 'Query' in infile.readline():
                            hf.skip_lines(infile, 3)
                        line = infile.readline()

    with open(outfilename, 'w', encoding='utf-8') as outfile:
        header = ('ID\tAverage_#_hits\tAverage_identity_%\t#_genomes_present_in'
                  '\t#_same_strand\tAverage_best_alignment_identity\n')
        outfile.write(header)
        for rna_id, value in srnas.items():
            value[1] /= value[0]
            value[0] /= value[2]
            value[3] /= value[0]
            value[4] /= value[2]
            outfile.write(rna_id + '\t')
            outfile.write(str(round(value[0], 1)) + '\t')
            outfile.write(str(round(value[1], 2)) + '\t')
            outfile.write(str(value[2]) + '\t')
            outfile.write(str(int(value[3])))
            # outfile.write(str(int(value[4])))
            outfile.write(str(value[4]))
            outfile.write('\n')


if __name__ == '__main__':
    main()
```
