## SupplementalFile2_python_scripts for "Assessing the conservation and targets of putative sRNAs in *Streptococcus pneumoniae*": srnarftarget_hits.py

```
"""
This is a script to read the csv output of
sRNARFTarget results and find the genes and
up/downstream genes in the GenBank files.
The output is in csv format so it can be
directly copied into excel
"""


import os
from typing import IO
import helper_functions as hf


def get_upstream(infile: IO[str]) -> str:
    """Get the upstream gene"""
    upstream: str = ''
    line: str = infile.readline()
    while '/product=' not in line:
        line = infile.readline()
    i: int = 31
    while line[i] != '"':
        if line[i] == '\n':
            line = infile.readline()
            i = 21
        upstream += line[i]
        i += 1
    return upstream


def get_downstream(infile: IO[str]) -> str:
    """Get the downstream gene"""
    downstream: str = ''
    line: str = infile.readline()
    while line[5:9] != 'gene':
        line = infile.readline()
    while '/product=' not in line:
        line = infile.readline()
    i: int = 31
    while line[i] != '"':
        if line[i] == '\n':
            line = infile.readline()
            i = 21
        downstream += line[i]
        i += 1
    return downstream


def get_target(infile: IO[str], mrna_id: str) -> tuple[str, str, str]:
    """Get the target gene"""
    infile.seek(0)
    prev: int = 0
    gene: str = ''
    while 'FEATURES' not in infile.readline():
        continue
    while True:
        line: str = infile.readline()
        if line[5:9] == 'gene':
            curr: int = infile.tell()
            line = infile.readline()
            i: int = 21
            ids: list[str] = []
            while line[i] == '/':
                ids.append(line)
                line = infile.readline()
            found: bool = False
            for rna_id in ids:
                if mrna_id in rna_id:
                    found = True
            if found:
                line = infile.readline()
                while '/product=' not in line:
                    line = infile.readline()
                i = 31
                while line[i] != '"':
                    if line[i] == '\n':
                        line = infile.readline()
                        i = 21
                    gene += line[i]
                    i += 1
                infile.seek(prev)
                upstream: str = get_upstream(infile)
                infile.seek(curr)
                downstream: str = get_downstream(infile)
                return (gene, upstream, downstream)
            prev = curr
        if not line:
            break
    return ('', '', '')


def main() -> None:
    """The main function"""
    path: str = os.getcwd()[:-7]
    srnarf: str = path + 'sRNARFTarget stuff/'
    directory: str = srnarf + 'Transcriptome100-300/'
    for direc in os.listdir(directory):
        if os.path.isdir(directory + direc):
            strain: str = direc.split(' ')[0]
            tsv: str = directory + direc + '/Top_hit.tsv'
            genbank: str = path + 'Annotated genomes/' + strain + '.gb'
            output: str = directory + direc + '/sRNARFTargetHits.tsv'
            with (open(tsv, 'r', encoding='utf-8') as infile1,
                  open(genbank, 'r', encoding='utf-8') as infile2,
                  open(output, 'w', encoding='utf-8') as outfile):

                hf.skip_lines(infile1)
                header: str = ('sRNA_ID\tmRNA_ID\tProbability\tGene\t'
                               'Upstream-gene\tDownstream-gene\n')
                outfile.write(header)
                num_srnas: int = 60
                for _ in range(num_srnas):
                    line: str = infile1.readline().strip()
                    srna_id: str = line.split(' ')[0]
                    mrna_id: str = line.split(' ')[1]
                    probability: str = line.split(' ')[2]
                    outfile.write(f'{srna_id}\t{mrna_id}\t{probability}\t')
                    gene: str = ''
                    upstream: str = ''
                    downstream: str = ''
                    gene, upstream, downstream = get_target(infile2, mrna_id)
                    outfile.write(f'{gene}\t{upstream}\t{downstream}\n')


if __name__ == '__main__':
    main()
```
