## SupplementalFile2_python_scripts for "Assessing the conservation and targets of putative sRNAs in *Streptococcus pneumoniae*": srnarftarget_top_5_hits.py

```
"""
Read the data output from sRNARFTarget
and write the top 5 hits for each sRNA
into a new file and the top 1 hit into a new file.
The output files will have the top 5 and 1 hits for
each sRNA.
"""


import os
import pandas as pd


def main() -> None:
    """The main function"""
    path: str = os.getcwd()[:-7]
    # destination: str = path + 'sRNARFTarget stuff/Transcriptome_only/'
    destination: str = path + 'sRNARFTarget stuff/Transcriptome100-300/'
    for directory in os.listdir(destination):
        if os.path.isdir(destination + directory):
            strain: str = directory.split(' ')[0]
            file: str = destination + directory + '/' + strain + '_results100-300.tsv'
            df = pd.read_csv(file, sep='\t')

            srnas: list[str] = []
            for _, row in df.iterrows():
                if row['sRNA_ID'] not in srnas:
                    srnas.append(row['sRNA_ID'])

            n = len(srnas)
            for i in range(n):
                swapped = False
                for j in range(0, n - i - 1):
                    if int(srnas[j][1:-1]) > int(srnas[j+1][1:-1]):
                        srnas[j], srnas[j+1] = srnas[j+1], srnas[j]
                        swapped = True
                if swapped is False:
                    break

            with open(destination + directory + '/' + 'Top_5_hits.tsv', 'w',
                      encoding='utf-8') as outfile:
                for column_name in df.columns:
                    outfile.write(column_name + '\t')
                outfile.write('\n')
                for srna_id in srnas:
                    output = df[df['sRNA_ID'] == srna_id].head(5) # type: ignore
                    outfile.write(output.to_string(index=False, header=False) + '\n')

            with open(destination + directory + '/' + 'Top_hit.tsv', 'w',
                      encoding='utf-8') as outfile:
                for column_name in df.columns:
                    outfile.write(column_name + '\t')
                outfile.write('\n')
                for srna_id in srnas:
                    output = df[df['sRNA_ID'] == srna_id].head(1) # type: ignore
                    outfile.write(output.to_string(index=False, header=False) + '\n')


if __name__ == '__main__':
    main()
```
