## SupplementalFile2_python_scripts for "Assessing the conservation and targets of putative sRNAs in *Streptococcus pneumoniae*": targetrna3_hits.py

```
"""
Find the upstream, downstream, and target length
of the predicted target
Output as a .csv so it can be copy pasted
into excel.
"""


import os
from typing import IO


def write_header(outfile: IO[str]) -> None:
    """Write the header for the output file"""
    outfile.write('sep=;\n')
    header: str = ('sRNA_ID;sRNA_subseq_start;sRNA_subseq_end;'
              'target;strand;target_length;target_start;target_end;mRNA_subseq_start;'
              'mRNA_subseq_end;energy;P-value;probability;annotation;upstream_gene;'
              'upstream_start;upstream_end;downstream_gene;downstream_start;'
              'downstream_end;interaction\n')
    outfile.write(header)


def write_data(data: tuple[str, ...], outfile: IO[str]) -> None:
    """Write the data for the output file"""
    for i, item in enumerate(data):
        if i != len(data) - 1:
            outfile.write(item + ';')
        else:
            outfile.write(item + '\n')


def get_upstream(infile: IO[str]) -> tuple[str, ...]:
    """Get the upstream gene"""
    upstream: str = ''
    line: str = infile.readline()
    while line[5:9] != 'gene':
        line = infile.readline()
    gene_start: str = ''
    gene_end: str = ''
    gene_start, gene_end = line[21:].split('..')
    if 'comp' in gene_start:
        gene_start = gene_start.split('(')[1]
        gene_end = gene_end.split(')')[0]
    if '>' in gene_end:
        gene_end = gene_end[1:]
    if '<' in gene_start:
        gene_start = gene_start[1:]
    line = infile.readline()
    while '/product=' not in line:
        line = infile.readline()
    i: int = 31
    while line[i] != '"':
        if line[i] == '\n':
            line = infile.readline()
            i = 21
            upstream += ' '
        upstream += line[i]
        i += 1
    return (upstream, gene_start, gene_end)


def get_downstream(infile: IO[str]) -> tuple[str, ...]:
    """Get the downstream gene"""
    downstream: str = ''
    line: str = infile.readline()
    while line[5:9] != 'gene':
        line = infile.readline()
    gene_start: str = ''
    gene_end: str = ''
    gene_start, gene_end = line[21:].split('..')
    if 'comp' in gene_start:
        gene_start = gene_start.split('(')[1]
        gene_end = gene_end.split(')')[0]
    if '>' in gene_end:
        gene_end = gene_end[1:]
    if '<' in gene_start:
        gene_start = gene_start[1:]
    line = infile.readline()
    while '/product=' not in line:
        line = infile.readline()
    i: int = 31
    while line[i] != '"':
        if line[i] == '\n':
            line = infile.readline()
            i = 21
            downstream += ' '
        downstream += line[i]
        i += 1
    return (downstream, gene_start, gene_end)


def get_target(infile: IO[str], target: str) -> tuple[str, ...]:
    """Get the target gene"""
    temp: str = ''
    if '(' in target:
        temp = target.split('(')[-1]
        target = temp[:-1]
    infile.seek(0)
    prev: int = 0
    while 'FEATURES' not in infile.readline():
        continue
    up: tuple[str, ...] = ()
    down: tuple[str, ...] = ()
    while True:
        line: str = infile.readline()
        strand: str = 'top'
        if line[5:9] == 'gene':
            curr: int = infile.tell()
            gene_start: str = ''
            gene_end: str = ''
            gene_start, gene_end = line[21:].split('..')
            if 'comp' in gene_start:
                strand = 'comp'
                gene_start = gene_start.split('(')[1]
                gene_end = gene_end.split(')')[0]
            if '>' in gene_end:
                gene_end = gene_end[1:]
            if '<' in gene_start:
                gene_start = gene_start[1:]
            line = infile.readline()
            i: int = 21
            ids: list[str] = []
            while line[i] == '/':
                ids.append(line)
                line = infile.readline()
            found: bool = False
            for srna_id in ids:
                if target in srna_id:
                    found = True
            if found:
                line = infile.readline()
                while '/product=' not in line:
                    line = infile.readline()
                i = 31
                while line[i] != '"':
                    if line[i] == '\n':
                        line = infile.readline()
                        i = 21
                    i += 1
                if prev - 100 >= 0:
                    infile.seek(prev - 100)
                    up = get_upstream(infile)
                else:
                    up = ('N/A', 'N/A', 'N/A')
                infile.seek(curr)
                down = get_downstream(infile)
                length = int(gene_end) - int(gene_start) + 1
                if strand == 'comp':
                    return (strand, str(length), gene_end, gene_start,
                            down[0], down[1], down[2],
                            up[0], up[1], up[2])
                return (strand, str(length), gene_start, gene_end,
                        up[0], up[1], up[2],
                        down[0], down[1], down[2])
            prev = curr
        if not line:
            break
    return ('', '', '', '', '', '', '', '', '', '')


def main() -> None:
    """The main function"""
    path: str = os.getcwd()[:-7]
    targetrna: str = path + 'TargetRNA3 stuff/'
    strains: list[str] = ['TIGR4',
               'D39',
               'TVO_1901920',
               'TVO_1902277',
               'TVO_Taiwan19F-14'] # Cannot do TVO_1901934 for GB version reasons
    for strain in strains:
        tsv: str = targetrna + strain + '_results/' + strain + '_summary.tsv'
        genbank: str = path + 'Annotated genomes/' + strain + '.gb'
        data: str = targetrna + strain + '_results/' + strain + '_updated.csv'
        with (open(tsv, 'r', encoding='utf-8') as infile1,
              open(genbank, 'r', encoding='utf-8') as infile2,
              open(data, 'w', encoding='utf-8') as outfile):

            write_header(outfile)
            line: str = infile1.readline().strip()
            while line:
                (srna_id, _, target, energy, p_value, probability, srna_start,
                 srna_stop, target_start, target_end, annotation, interaction) = line.split('\t')
                (strand, target_length, gene_start, gene_end,
                 upstream_gene, up_gene_start, up_gene_end,
                 downstream_gene, down_gene_start, down_gene_end) = get_target(infile2, target)
                info: tuple[str, ...] = (srna_id, srna_start, srna_stop, target, strand,
                        target_length, gene_start, gene_end,
                        target_start, target_end, energy, p_value, probability, annotation,
                        upstream_gene, up_gene_start, up_gene_end,
                        downstream_gene, down_gene_start, down_gene_end, interaction)
                write_data(info, outfile)
                line = infile1.readline().strip()


if __name__ == '__main__':
    main()
```
