## SupplementalFile2_python_scripts for "Assessing the conservation and targets of putative sRNAs in *Streptococcus pneumoniae*": targetrna3_top_hits.py

```
"""
Organize the top hit of each sRNA for
TargetRNA3 into a single .tsv that can
be easily copy/pasted into excel
"""


import os
import helper_functions as hf


def main() -> None:
    """The main function"""
    path: str = os.getcwd()[:-7]
    strain: str = 'suis'
    results: str = path + 'pyogenes mutans suis/' + strain + '_results/'
    outfilename: str = results + strain + '_summary.tsv'

    srnas: dict[str, str] = {}

    for filename in os.listdir(results):
        if filename[-4:] == '.txt':
            infilename: str = results + filename
            with open(infilename, 'r', encoding='utf-8') as infile:
                name: str = filename.split('_')[0]
                hf.skip_lines(infile)
                line: str = infile.readline()
                srnas[name] = line

    with open(outfilename, 'w', encoding='utf-8') as outfile:
        # 81 for the highest number identifier for any of the sRNAs (M81)
        # This is for sorting them from M1 -> M2 -> ... -> M81
        for i in range(82):
            name = 'M' + str(i)
            if name in srnas:
                outfile.write(f'{name}\t{srnas[name]}')


if __name__ == '__main__':
    main()
```
