## SupplementalFile2_python_scripts for "Assessing the conservation and targets of putative sRNAs in *Streptococcus pneumoniae*": transcriptome.py

```
"""
This is a script to make a transcriptome fasta
file using the genome and annotated genome files.
The output file will have all the mRNAs that
serve as input for any of the targer prediction
programs.
"""


import os
import helper_functions as hf


NUMS = '0123456789'

def main() -> None:
    """The main function"""
    path: str = os.getcwd()[:-7]
    fastas: str = path + 'TVO fasta files/'
    genbanks: str = path + 'Annotated genomes/'
    trans: str = path + 'Transcriptomes100-300/'
    strain: str = 'TVO_1902277'
    file1: str = fastas + strain + '.fasta'
    file2: str = genbanks + strain + '.gb'
    file3: str = trans + strain + '_transcriptome100-300.fasta'
    with (open(file1, 'r', encoding='utf-8') as infile1,
          open(file2, 'r', encoding='utf-8') as infile2,
          open(file3, 'r', encoding='utf-8') as outfile):

        count: int = 0
        stop: int = 0
        while infile2.readline()[:8] != 'FEATURES':
            continue
        while True:
            if stop:
                break
            line: str = infile2.readline()
            if line[:6] == 'ORIGIN' or not line:
                break
            if line[5:9] == 'gene':
                # print('Gene:', count)
                # upstream, downstream = 0, 0
                upstream: int = 100
                downstream: int = 300
                count += 1
                i: int = 0
                while line[i] not in NUMS:
                    i += 1
                start: str = ''
                end: str = ''
                start, end = line[i:].strip().split('..')
                if '<' in start:
                    start = start[1:]
                if '>' in end:
                    end = end[1:]
                strand: str = '+'
                if 'complement' in line:
                    strand = '-'
                    end = end[:-1]
                start_int = 0
                end_int = 0
                start_int, end_int = int(start), int(end)
                line = infile2.readline()
                gene: str = ''
                new_locus: str = ''
                old_locus: str = ''
                if 'gene' in line:
                    gene = line.split('"')[1]
                    new_locus = infile2.readline().split('"')[1]
                    try:
                        old_locus = infile2.readline().split('"')[1]
                        outfile.write(f'>{gene} ({old_locus})\n')
                    except ValueError:
                        outfile.write(f'>{gene} ({new_locus})\n')
                else:
                    new_locus = line.split('"')[1]
                    line = infile2.readline()
                    try:
                        old_locus = line.split('"')[1]
                        outfile.write(f'>{old_locus}\n')
                    except ValueError:
                        outfile.write(f'>{new_locus}\n')
                return_to: int = infile2.tell()
                line = infile2.readline()
                while line[5:9] != 'gene':
                    line = infile2.readline()
                    if 'ORIGIN' in line:
                        stop = 1
                        break
                next_start: str = ''
                j: int = 0
                while j < len(line) and line[j] not in NUMS:
                    j += 1
                if j != len(line):
                    while line[j] in NUMS:
                        next_start += line[j]
                        j += 1
                    next_start_int: int = int(next_start)
                    if next_start_int - end_int < downstream:
                        downstream = next_start - end # type: ignore
                    infile2.seek(return_to)
                sequence = hf.retrieve_sequence(start_int, end_int, infile1,
                                                upstream, downstream) # type: ignore
                if strand == '-':
                    sequence = hf.reverse_complement(sequence)
                outfile.write(sequence + '\n\n')


if __name__ == '__main__':
    main()
```
